# Streptococcal meningoencephalitis dynamically shapes developmental monocyte and macrophage trajectories in the CNS

**DOI:** 10.1101/2024.02.27.582183

**Authors:** Vitka Gres, Florens Lohrmann, Vidmantė Fuchs, Jana Neuber, Zohreh Mansoori Moghadam, Anne K. Lösslein, Lance F. P. Bosch, Tiago Figueiral-Martins, Sebastian Baasch, David Obwegs, Gianni Monaco, Julia Henschel, Klaus P. Knobeloch, Marco Prinz, Steffen Jung, Katrin Kierdorf, Sagar, Julia Kolter, Daniel Erny, Philipp Henneke

## Abstract

Macrophages in the meninges, especially in the dura mater sheathing the brain from the skull, are involved in the immune defense of the central nervous system (CNS). However, their site-specific origin and function, both in steady state and in bacterial CNS infections are incompletely understood. Using an intravenous model of streptococcal meningoencephalitis that mimics hematogenous dissemination in humans, we found that bacteria accumulated predominantly in the leptomeninges and dura, whereas invasion into the brain parenchyma was rare. However, monocyte infiltration into the leptomeninges and parenchyma strongly correlated with disease severity.

In the dura, infection triggered activation and loss of resident macrophages, followed by rapid engraftment of inflammatory monocytes that transiently replenished the dural macrophage niche. Under homeostasis, dural monocytes were supplied independently of CCR2 from adjacent skull bone marrow. In infection, however, this local reservoir was rapidly exhausted, and the markedly increased demand for monocytes required mobilization from peripheral bone marrow sources, revealing context-dependent heterogeneity in monocyte origin. Infection also reshaped ontogeny of this differential monocyte output, with an increase in Monocyte-Dendritic Cell Progenitor - derived monocytes (MDP-Mo). MDP-Mo exhibited enhanced MHC-II expression and persisted in the brain during the resolution phase together with CD4⁺ T cells, suggesting a role in antigen presentation after bacterial clearance.

Together, these findings reveal a highly dynamic and compartment-specific remodeling of monocyte ontogeny, recruitment, and differentiation across CNS borders during bacterial meningoencephalitis. These mechanisms may offer opportunities for therapeutic interventions in the future.

**One Sentence Summary:** Streptococcal meningoencephalitis disrupts homeostatic, skull bone marrow-derived monocyte and macrophage trajectories in the dura.

## 2. INTRODUCTION

Bacterial meningoencephalitis is a devastating disease that affects millions of people each year. Group B streptococcus (*Streptococcus agalactiae*, GBS) is a common colonizer of the gastrointestinal and urogenital tract, but also a leading cause of neonatal meningoencephalitis. GBS poses a particular risk to patients with diabetes mellitus, pregnant women, the elderly, and immunocompromised people (*1*). GBS meningoencephalitis has a mortality rate of up to 10%, and surviving patients often suffer from neurological sequelae (*2*, *3*).

The integrity of the brain is protected by anatomical barriers that ensure the controlled passage of molecules from the periphery into the central nervous system (CNS). The choroid plexus, the perivascular space along the vessels, and the meninges harbor populations of macrophages (Macs), together called CNS-associated macrophages (CAM). They play a crucial role in maintaining homeostasis of the CNS, but also provide neuroprotection in many pathologies (*4–7*). Anatomically, the meninges consist of three layers – *dura mater*, as well as *arachnoid* and *pia mater* (leptomeninges) (*8*). The dura harbors a complex immunologic landscape that includes T-and B-cells, natural killer cells, neutrophils, plasmacytoid dendritic cells (pDCs), conventional DCs (cDCs), mast cells, monocytes, and diverse subsets of the most abundant population – dural macrophages (DM) (*9*).

Dural sinuses, which drain venous blood, are sites of antigen sampling and presentation (*10*). The dura is connected to the skull marrow by ossified channels, which provide a direct migration route for monocytes, neutrophils and B-cells (*11*, *12*). Recently, cerebrospinal fluid has been described to transport bacteria from the dura to the skull marrow when bacteria like *Streptococcus pneumoniae* were directly injected into the cisterna magna (*13*). However, the dynamics of the dura–skull marrow immune axis in blood-borne bacterial infection remain unclear.

In contrast to the dura, the brain parenchyma contains only one resident immune population in steady state – microglia (MG). In a healthy brain, MG are highly unlikely to encounter peripheral immune cells such as neutrophils, T cells or monocytes. In contrast, neuroinflammatory diseases, e.g., multiple sclerosis (*14*) or ischemic stroke (*15*) can lead to leakage of the blood-brain barrier and infiltration of peripheral leukocytes, contributing to MG activation and disease exacerbation.

During GBS infection in the CNS, neuroimmune communication appears to be crucial. On the one hand, MG are thought to exacerbate immunopathology in infection, e.g., by producing nitric oxide, which ultimately leads to neuronal death (*16*). On the other hand, nociceptors of the trigeminal ganglia in the dura dampen the response of DM through CGRP-RAMP1 signaling, which leads to increased spread of bacteria into the brain parenchyma (*17*). Hence, the cross-talk of the immune compartments in the CNS parenchyma and interfaces seem to play major roles in disease progression and outcome. However, a deeper understanding of the precise immune cell dynamics in the CNS and particularly its borders during streptococcal meningoencephalitis is still lacking.

Here, we investigated the discrete immune responses of MG, DM, and infiltrating monocytes in streptococcal meningoencephalitis caused by hematogenous spread. We took an integrated approach to understand the intersectionality of myeloid fates in infection, focusing on localization and differentiation trajectories. This allowed us to demonstrate that DM are positioned at the primary interface of the infection, initiating immune activation. Infiltrating Ly6C^hi^ monocytes actively replenished resident DM and were imprinted by their cellular origin. Moreover, monocyte invasion into the adjacent CNS parenchyma was associated with adverse infection outcome. Thus, streptococcal infection induces fundamental ontogenetic and functional changes in tissue-resident macrophages of the CNS interfaces, and the recruitment pathways and differentiation dynamics of migrating monocytes are linked to disease advancement.

## 3. RESULTS

### 3.1 Hematogenous GBS infection highlights the meninges as a reactive CNS barrier

Previous studies on the immune responses in bacterial meningoencephalitis in mice typically relied on disease models in which the pathogen was directly injected into the *cisterna magna* (*13*, *18*, *19*), therefore circumventing the effects of peripheral immune activation and systemic bacterial dissemination on CNS inflammation. In streptococcal meningoencephalitis, mucosal colonization usually precedes CNS infection, and bacterial dissemination via the blood is often observed (*20*). To investigate the systemic dissemination of GBS and its capacity to invade the CNS, we intravenously injected wild type (*Wt*) mice with 5x10^7^ CFU of the serotype III hypervirulent BM110 GBS strain, which is linked to neonatal meningoencephalitis in humans (*21*, *22*). The infection spread rapidly and the overall fitness of the infected mice drastically declined until 3 days post infection (dpi) when mice lost around 20% of their body weight, leading to termination of the experiments (Fig. S1A). We therefore focused on 2 dpi timepoint, at which we observed bacterial presence in blood, brain lysates including leptomeninges, dura mater, liver, skull and femur (Fig 1A). In agreement with previous data on GBS sepsis (*23*), interleukin (IL)-6 levels in the plasma (Fig. S1B) as well as tumor necrosis factor (TNF), IL-1β and interferon (IFN)-β in the brain lysates of infected mice were substantially increased (Fig. S1C).

**Fig. 1:**
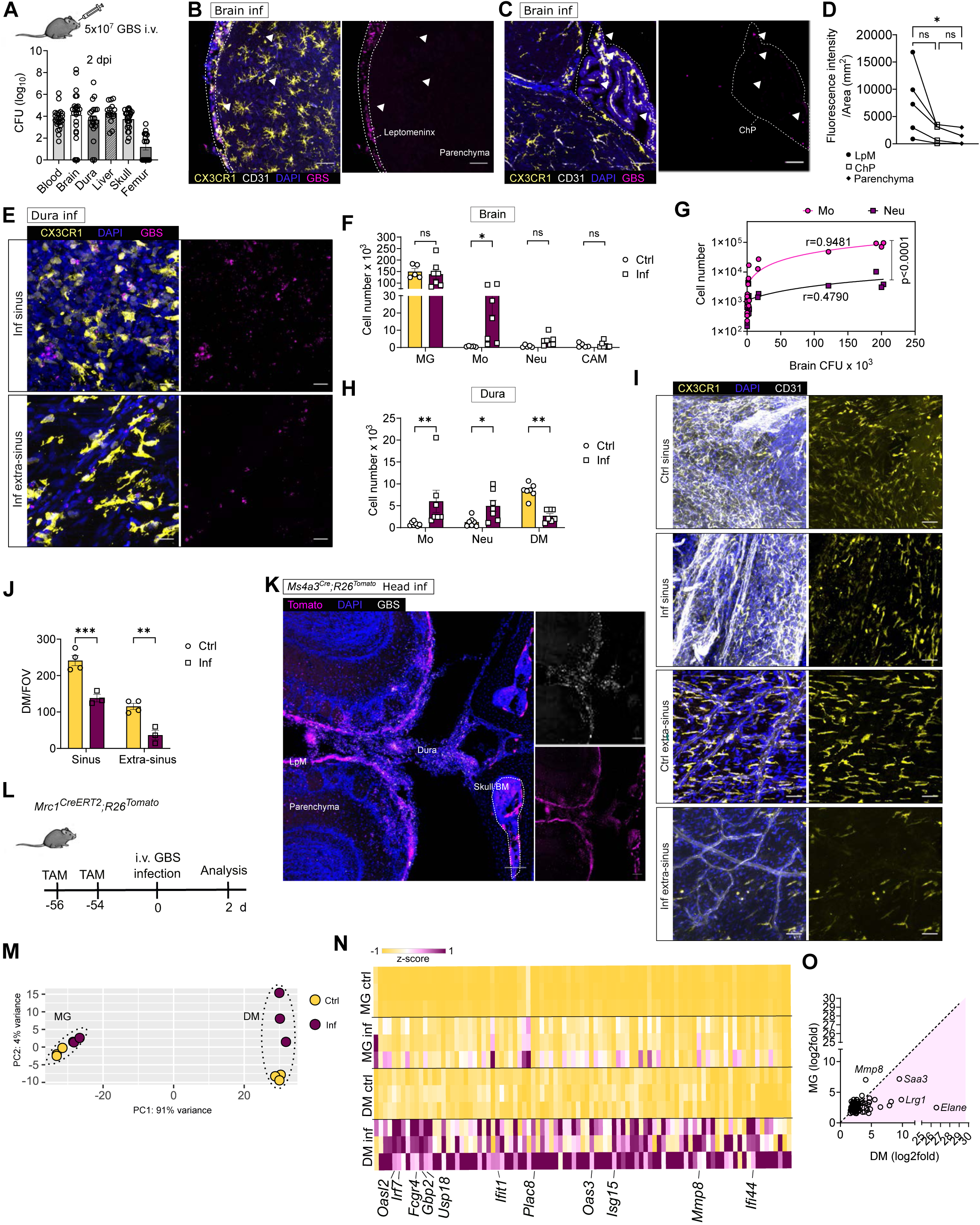
Hematogenous GBS infection highlights the meninges as a reactive CNS barrier. **A:** *Wt* mice were intravenously injected with 5x10^7^ CFU of the serotype III hypervirulent BM110 GBS strain. Colony forming units (CFU) counts from homogenized brains (n=25), dura (n=21), liver (n=16), extracted bone marrow from femur (n=19) and skull (n=25) (per tissue/organ) and blood culture (per ml) (n=24) at 2 dpi. Data represent at least five independent experiments. **B-C:** Representative maximal intensity projection of sagittal brain slices of infected *Cx3cr1^gfp/+^* mice including leptomeningeal area and parenchyma (**B**) and choroid plexus (ChP) (**C**) labelled with GBS-AF633 (magenta) and DAPI (blue). White arrows mark some of the GBS found in the parenchyma and choroid plexus. Scale bar = 50 µm. **D:** Fluorescence intensity of GBS-AF633 labelling divided by the area of leptomeninges (LpM), cortical parenchyma, or choroid plexus (ChP). For ChP, either the lateral or the third ventricle was imaged. Each dot represents an individual infected mouse, n=5. Statistics by paired t-test. *p<0.05, data represents at least three independent experiments. **E:** Representative maximal intensity projection of dural whole mount from infected *Cx3cr1^gfp/+^* mice labelled with GBS-AF633 (magenta) and DAPI (blue), from either region of the sinus or the extra-sinus region. Scale bar = 20 µm. **F:** Absolute numbers of microglia (MG), monocytes (Mo), neutrophils (Neu) and CNS-associated macrophages (CAM), including leptomeningeal, perivascular and choroid plexus Macs from control (Ctrl) and infected (Inf) mice measured by flow cytometry. One brain hemisphere per mouse was used for quantification. n=5-7, data represents at least two independent experiments. Multiple Mann-Whitney test followed by Holm-Šídák, *p<0.05, ns = not significant **G**: Correlation of CFU counts and total monocyte or neutrophil number in the brain assessed by Spearman correlation. Slopes compared by simple linear regression, n=20. Data represent at least three independent experiments. R represents the goodness of fit in simple linear regression. p<0.0001 **H:** Absolute numbers of monocytes, neutrophils and DM from control and infected dura mater measured by flow cytometry, n=7. Statistics by Multiple Mann-Whitney test followed by Holm-Šídák. Data represent at least two independent experiments. *p<0.05, **p <0.01. **I:** Representative maximal intensity projection of dural whole mount from control and infected *Cx3cr1^gfp/+^* mice stained with DAPI (blue) and labelled with CD31-PE (white) showing loss of GFP-positive DM, in sinus and extra-sinus area. Scale bar = 50 µm. **J:** Quantification of *Cx3cr1^gfp/+^* DM per field of view (FOV) in sinus and extra-sinus area of control and infected dura. 2-3 sinus and extra-sinus positions were quantified per mouse, and average values were plotted. N= 3-4, from two independent experiments. Statistics by two-way ANOVA, followed by Tukey’s multiple comparisons test. **p<0.01, ***p <0.001 **K:** Representative image of a coronal slice of infected *Ms4a3^Cre^;R26^Tomato^* mouse head, stained with DAPI (blue), and labelled with GBS-AF633 (white). Scale bar = 100 µm. **L:** Scheme of experimental time course. *Mrc1^CreERT2^;R26^Tomato^*mice were treated with tamoxifen twice, 8 weeks prior to the infection. At 2 dpi, DM and MG were sorted from dura and brain. Two mice were pooled for n=1, in total 6 mice were used (n=3). **M:** Principal component analysis (PCA) of bulk RNA-seq from sorted populations. **N:** Z-score heat map of 93 common upregulated DEG in both MG and DM, with log_2_(fold change) larger than 1.5 and padj<0.05. **O:** A comparison scatter plot of genes upregulated both in MG and DM, shown as a log_2_(fold change) of common upregulated DEG from MG and DM.

Detailed spatial mapping revealed that GBS dissemination into the brain parenchyma was rare, indicating that CNS border tissues effectively restricted bacterial entry (Fig. 1B, S1D). While bacteria were present in the choroid plexus at low levels (Fig. 1C, D, S1E), they accumulated predominantly at the leptomeninges and the dura mater (Fig. 1B, E, S1D, S1F). This containment implied active immune responses, prompting us to next examine immune cell recruitment in response to GBS invasion. Flow cytometry analysis showed a significant influx of Ly6C^hi^ monocytes into the brain, including the leptomeninges, while neutrophil numbers were only marginally increased (Fig.1F, S1G). Monocyte influx positively correlated with the bacterial burden in the brain (including leptomeninges) (r > 0.9), and in the mice with severe infections, monocyte counts increased more than 50-fold compared to less affected mice (Fig.1G). In contrast, the dura of GBS infected mice showed an increase in both monocyte and neutrophil counts (Fig.1H). Moreover, resident DM numbers in infection were significantly reduced, both within the sinuses and in extra-sinus area (Fig.1I, J). While DM have been previously characterized as MHC-II^hi^ or MHC-II^low^, both subsets declined proportionally during infection, with unchanged subset frequencies (Fig.S1H, I).

To spatially validate both GBS localization and bone marrow-derived myeloid cell influx, we utilized *Ms4a3^Cre^;R26^Tomato^*mice, in which granulocyte-monocyte progenitors (GMP) and their progeny (including monocytes and neutrophils) are labeled. Coronal head sections from infected *Ms4a3^Cre^;R26^Tomato^* mice showed robust accumulation of Tomato^+^ myeloid cells in both dura and leptomeninges, co-localizing with GBS foci (Fig.1K, S1J).

Collectively, these data show a compartmentalized pattern of GBS dissemination drastically affecting the immune cell composition of the meninges and the brain. Considering that tissue resident macrophages are the first responders in infection, we sought to investigate the transcriptional changes of CNS resident macrophage populations upon the bacterial challenge. Specifically, we focused on two distinct myeloid populations: DM and MG. To selectively label long-lived DM, we employed *Mrc1^CreERT2^;R26^Tomato^*mice with tamoxifen-induced labeling performed eight weeks prior to infection (Fig.1L). This timeline enabled us on one hand to distinguish resident DM from newly recruited monocyte-derived cells in the dura, while allowing us to discriminate between MG and other CAM on the other (Fig.S1K, L).

Principal component analysis (PCA) of the bulk RNA seq showed distinct clustering of the isolated populations. Whereas MG from control and infected mice clustered closely together, DM from infected and uninfected mice showed a distinct gene expression profile (Fig 1M). In line with our previous data on peripheral GBS infection (*24*) many of the upregulated differentially expressed genes (DEG) in DM and MG after GBS infection were associated with type I IFN signaling and with the defense response to bacteria, e.g., *Oasl2, Irf7, Usp18, Fcgr4, Gbp2, Ifit1, Oas3, Isg15,* and *Ifi44,* implying the existence of a common type I IFN-driven response of tissue resident macrophages to GBS (Fig.1N). While both populations mounted a similar core transcriptional response, the overall magnitude of gene induction expressed as log_2_(fold change) was higher in DM compared to MG (Fig.1O).

In summary, a strong inflammatory response and reduction of DM, coupled with a monocyte and neutrophil influx establishes the dura as a highly primary reactive immune interface in response to GBS infection.

### 3.2 Loss of dural macrophages triggers compensatory monocyte differentiation

Previous reports have shown that neuroinflammatory conditions such as infection or stroke enable monocyte engraftment into both the parenchyma and in the meninges (*18*, *25–27*). We therefore asked whether the changes in resident DM and MG described above were accompanied by high infection-driven monocyte-to-macrophage differentiation. As a baseline, we first examined monocyte-macrophage turnover in homeostasis using the *Cxcr4^CreERT2^;R26^Tomato^* mouse model, in which tamoxifen (TAM) induction labels hematopoietic stem cell (HCS)-derived cells, for up to six months (*28*). In steady state, MG are established early in embryonic development and do not receive substantial input from peripheral monocytes (*29*, *30*) and thus remained unlabeled throughout the experiment. In contrast, DM showed a gradual replacement by circulating monocytes, reaching up to 24% after 8 weeks and 45% at 16 weeks (Fig. S2A). Next, we assessed whether the infection altered DM and MG differentiation dynamics and analyzed *Cxcr4^CreERT2^;R26^Tomato^* mice 2 dpi that were treated with TAM 8 weeks prior (Fig. 2A). We did not detect Tomato^+^ MG in the brain parenchyma or an increased frequency of Tomato^+^ DM in infection (Fig. 2B). In the dura, however, GBS infection induced a marked expansion of late-Mo/early-Mac cells co-expressing Ly6C and CD206 (Fig. 2C, D). These Macs were of HSC origin (Fig. 2E) and, unlike Ly6C⁺ dural monocytes, upregulated MHC-II (Fig. 2F). Therefore, this population likely represented a transient monocyte-to-Mac differentiation stage that facilitates repopulation of the DM compartment during infection, compensating for infection-driven DM loss.

**Fig. 2:**
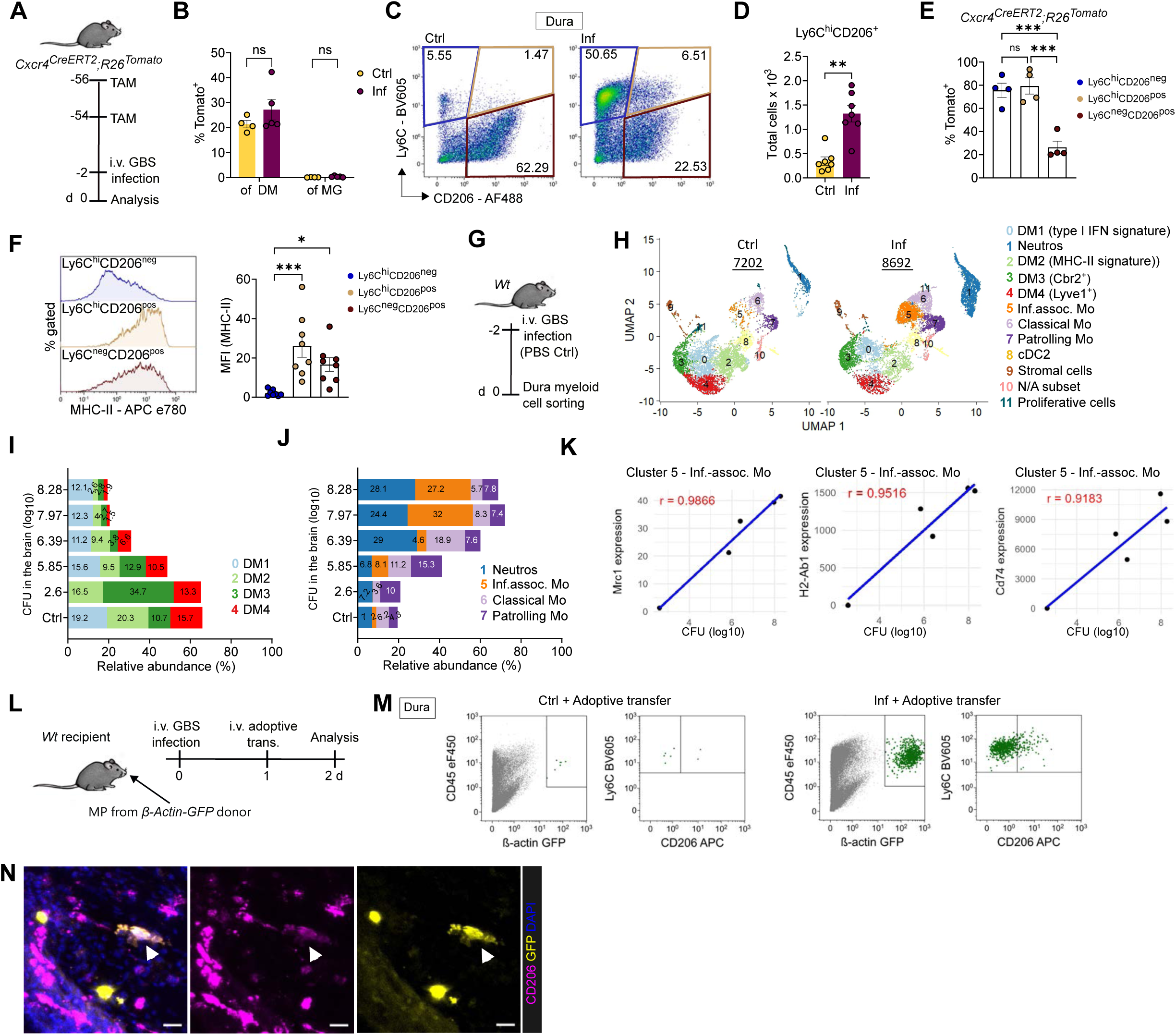
Loss of dural macrophages triggers compensatory monocyte differentiation. **A:** Scheme of experimental timeline. *Cxcr4^CreERT2;^R26^Tomato^* mice were induced with tamoxifen (TAM) 8 weeks prior to GBS infection and sacrificed at 2 dpi. **B:** Percentages of Tomato-labelled CD45^hi^CD11b^+^Ly6C^neg^Ly6G^neg^CD206^+^ DM in dura and CD45^lo^CD11b^+^Ly6C^neg^Ly6G^neg^CD206^neg^ MG in brain. Statistic by Mann-Whitney U-test n=4-5. Data represent three independent experiments. ns=not significant. **C:** Representative flow cytometry plots showing a presence of Ly6C^hi^CD206^+^ population in infected and control mice. Numbers in each gate indicate the percentage of the population. **D:** Flow cytometry-based quantification of Ly6C^hi^CD206^+^ population in Ctrl and Inf mice, n=7 Statistics by Mann-Whitney U-test. Data represent three independent experiments. **p<0.01 **E:** Percentage of Tomato-labeling in *Cxcr4^CreERT2;^R26^Tom^* mice in Ly6C^hi^CD206^+^ population in comparison to Ly6C^hi^CD206^neg^ monocytes and Ly6C^neg^CD206^+^ DM in infected dura. n=4, representing two independent experiments. Statistics by ordinary one-way ANOVA, followed by Dunnett’s multiple comparisons test. ns= not significant, ***p<0.001 **F:** MHC-II expression in Ly6C^hi^CD206^+^ population in comparison to Ly6C^hi^CD206^neg^ monocytes and Ly6C^neg^CD206^+^ DM. MHC-II expression is visualized as mean intensity fluorescence (MFI). n=8, representing three independent experiments. Statistics by ordinary one-way ANOVA, followed by Tukey’s multiple comparisons test. *p<0.05, ***p<0.001. **G:** Scheme of experimental timeline used for sorting of myeloid population for single cell sequencing. **H:** Uniform Manifold Approximation and Projection (UMAP) representation based on gene expression profiling of sorted CD45^+^CD11b^+^ myeloid population from the control and infected dura of *Wt* mice. The number of analyzed cells per condition is underlined. **I:** Relative abundance of DM clusters (0, 2, 3, 4) expressed as percentages of total myeloid cells, according to the individual bacterial burden in the brain, or the overall percentages from control mice. Each color represents a specific cluster indicated in **H**. **J:** Relative abundance of neutrophil and monocyte clusters (1, 5, 6, 7) expressed as percentages of total myeloid cells, according to the individual bacterial burden in the brain, or the overall percentages from control mice. Each color represents a specific cluster indicated in **H**. **K**: Linear regression of *Mrc1*, *H2-Ab1* and *Cd74* expression in infection-associated monocytes (cluster 5) against GBS CFU counts in the brain. To calculate R, Pearson correlation coefficient was used between its average expression level and the CFU count of the respective replicate. **L:** Scheme of experimental setup for adoptive transfer experiment. *Wt* mice were infected and 1 dpi received 2-4x10^5^ monocyte progenitors (MP) sorted from bone marrows *of β-Actin-GFP* mice. Control mice were administered with PBS before receiving transferred monocyte progenitors. **M:** Flow cytometry plots showing partial upregulation of CD206 in donor *β-Actin-GFP* mice bone marrow monocyte progenitors transferred to infected *Wt* mice. **N:** Representative image of infected dura, a *β-Actin-GFP* mice bone marrow donor-derived monocyte progenitor upregulating CD206 is marked with arrow. Scale bar = 20 µm.

To further explore changes of myeloid populations in depth, we performed single-cell transcriptional analysis on FACS sorted CD45^+^CD11b^+^ myeloid cells from control and GBS infected dura (Fig. 2G). Using unbiased hierarchical clustering analysis, we were able to distinguish 12 distinct cell clusters (Fig. 2H, S2B). Within the Mac compartment, all DM clusters expressed canonical DM genes, including *Mrc1*, *Pf4*, *Cd68*, and *Cx3cr1*, but each exhibited some distinct transcriptional features (Fig. S2B). Cluster 0 was enriched for type I IFN–responsive genes such as *Oasl1* and *Cxcl10*. Cluster 2 showed elevated expression of MHC-II–associated genes, consistent with enhanced antigen-presenting potential. Cluster 3 was marked by higher *Cbr2* expression, while cluster 4 was characterized by increased *Lyve1* (Fig. S2C). Furthermore, we identified three monocyte populations: classical monocytes, patrolling monocytes characterized by higher *Itgal* expression, and a distinct population that emerged specifically during infection and was virtually absent under homeostatic conditions (cluster 5) (Fig. 5H, S2B). We therefore refer to this population as infection-associated monocytes, which exhibited elevated expression of antimicrobial and inflammatory genes, including *Nos2, Rsad2, Mif, Cfb, Cxcl9,* and *Ccl5,* compared to the other monocyte clusters (Fig. S2D).

Consistent with previous observations, a relative decrease of percentage in macrophage clusters (0,2,3,4, Fig. 2I) and an increase of percentage in neutrophil (1) and monocyte (5,6,7) clusters (Fig. 2J) corelated to the bacterial burden in the brain (Fig. S2E). DM, particularly clusters 0 and 3, upregulated necroptosis regulators *Mlkl, Ripk1* and *Zbp1*, indicating that their loss may involve necroptotic death pathways reported in other bacterial infections (Fig. S2F)(*31*, *32*). Conversely, the expression of *Mrc1*, *H2-Ab1* and *Cd74* in monocyte cluster 5 increased proportionally to the bacterial burden (Fig. 2K), consistent with accelerated monocyte-to-macrophage differentiation in response to infection-driven macrophage depletion (Fig. 2C–F).

Next, to functionally validate the rapid monocyte-to-macrophage differentiation observed during infection, we adoptively transferred monocyte progenitors (MP) isolated from *β-actin-GFP* mice into either infected or control mice (Fig. 2L, S2G). In infected mice, we detected robust engraftment of GFP⁺ donor cells in the dura, with preferential accumulation in sinus-associated regions (Fig. S2H). A subset of these cells upregulated CD206 and acquired an ameboid morphology, consistent with acquisition of macrophage-like features (Fig. 2M, N). Overall, infection triggered a broad diversification of monocyte states and accelerated their differentiation into Macs, highlighting a remarkable degree of lineage plasticity within the inflamed dura.

### 3.3 Distinct monocyte sources support dural immunity in steady state and infection

Having established that infection induces extensive monocyte recruitment into the dura and the brain, we next wanted to determine their functional role in streptococcal infection. Since CCR2 is considered a key driver of monocyte mobilization from the bone marrow, we performed GBS infection in *Ccr2^-/-^*mice.

At 2 dpi, bacterial loads were similar between GBS-infected *Wt* and *Ccr2^-/-^* mice in the periphery and the brain, including leptomeninges (Fig. S3A), indicating that infiltrating Ly6C^hi^ monocytes neither play a critical role in early bacterial control, nor were directly involved in bacterial dissemination into the CNS. Notably, steady state monocyte numbers in the dura were similar between *Ccr2^-/-^* and *Wt* mice (Fig. 3A), consistent with the observation that dural monocytes express significantly lower levels of *Ccr2* than circulating monocytes (Fig. S3B). In contrast, we did not observe a significant monocyte influx in *Ccr2^-/-^* mice in the dura or in the brain upon infection, compared to the situation in *Wt* mice (Fig. 3A, 3B). Lack of monocytes was not compensated for by higher numbers of infiltrating granulocytes during GBS infection, as *Ccr2^-/-^* mice showed even lower dural neutrophil numbers than *Wt* mice (Fig. S3C).

**Fig. 3:**
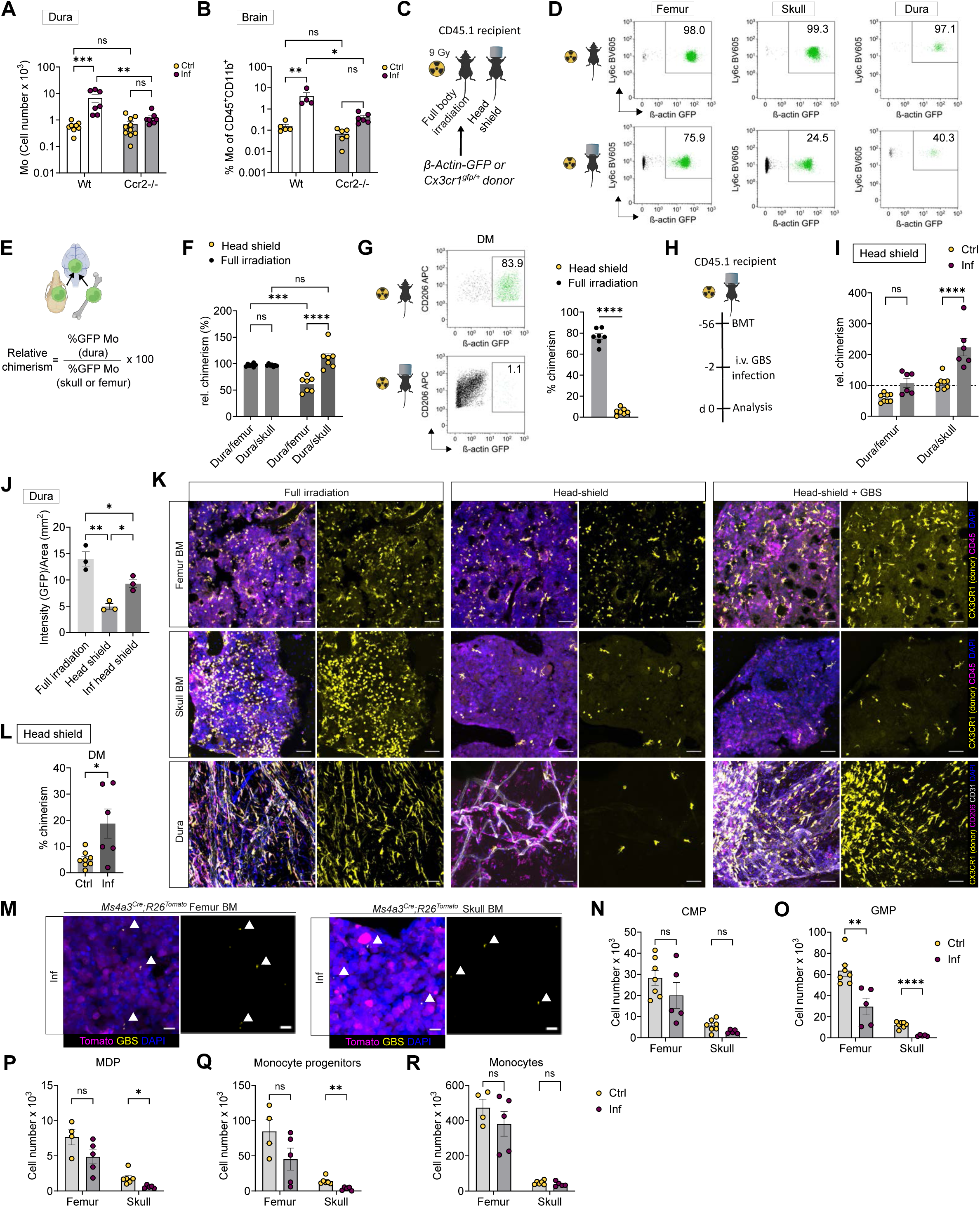
Distinct monocyte sources support dural immunity in steady state and infection. **A:** Flow cytometry quantitative analysis of monocytes from dura of control and infected *Wt* or *Ccr2^-/-^* mice. n=7-10, from at least three independent experiments. Statistics by two-way ANOVA and Tukey’s multiple comparisons test **p<0.01, ***p<0.001, ns=not significant. **B:** Flow cytometry analysis of monocyte frequencies from the brains of control and infected *Wt* or *Ccr2^-/-^* mice. n=4-6, from at least two independent experiments. Statistics by two-way ANOVA and Tukey’s multiple comparisons test, *p<0.05, **p<0.01, ns=not significant. **C:** Scheme of different strategies of radiation procedure followed by bone marrow transplantation from *Cx3cr1^+/gfp^* or *β-Actin-GFP* mice. Analysis was done 8 weeks post bone marrow transfer. **D:** Representative flow cytometry plots depicting donor-originating monocytes in green, from either fully irradiated mice or head-shielded mice. Numbers in gates represent % of gated GFP-positive (donor-derived) monocytes. **E:** Scheme of calculation of relative chimerism with formula used. Illustration made in BioRender (biorender.com). Monocytes= Mo. **F:** Chimerism rates of monocytes in dura compared to chimerism of skull monocytes, or femur monocytes. Statistics by two-way ANOVA followed by Tukey’s multiple comparisons test. n=7, data represent at least three independent experiments. ***p<0.001, ****p<0.0001, ns=not significant. **G:** On the left, representative flow cytometry plots depicting donor-originating DM in green, from either fully irradiated mice or head-shielded mice. Numbers in gates represent % of gated GFP-positive (donor-derived) DM. On the right, chimerism rates of DM in fully irradiated or head-shielded mice. Statistics by Mann-Whitney U – test. n=7, data represent three independent experiments. ****p<0.0001. **H:** Scheme of experimental timeline post head-shielded irradiation. 8 weeks post head-shielded irradiation and bone marrow transfer (BTM), mice were infected with GBS and analyzed 2 dpi. **I:** Chimerism rates of monocytes in dura compared to chimerism of skull monocytes, or femur monocytes, in control or infected head shielded mice. n=6-8 from at least three independent experiments. Statistics by two-way ANOVA followed Šídák’s multiple comparisons test, ****p<0.0001, ns = non significant. **J:** Fluorescent intensity of GFP signal in divided by area in whole mounted dura as shown in representative images in S3I. N=3, *p<0.05, **p<0.01 **K:** Representative microscopy images of femur or skull bone marrow, or whole mount dura after bone marrow transfer from *Cx3cr1^gfp/+^* donor. The mice were either fully irradiated, or they were head shielded. A group of head shielded mice was infected. Scale bar = 50 µm. **L:** Chimerism of DM post infection measured by flow cytometry. Statistics by unpaired t-test. n=6-8, from at least three independent experiments. **\***p< 0.05 **M:** Femur and skull bone marrow from infected *Ms4a3^Cre^;R26^Tomato^* mice labelled with GBS-AF633 (depicted in yellow). GBS is marked with white arrows. Scale bar =10 µm. **N:** Numbers of CMP per femur or skull marrow in steady state and infection. Statistics by multiple unpaired t-test followed by the Holm-Šídák, n=5-7, data represents at least two independent experiments. ns = not significant. **O:** Numbers of GMP of femur or skull marrow in steady state and infection. Statistics by multiple unpaired t-test followed by the Holm-Šídák, n=5-7, data represents at least two independent experiments. ***p < 0.001, ****p < 0.0001. **P:** Numbers of MDPs of femur or skull marrow in steady state and infection. Statistics by multiple unpaired t-test followed by the Holm-Šídák, n=4-6, data represents at least two independent experiments. ns = not significant, ***p < 0.001. **Q:** Numbers of monocyte progenitors (including cMOP and iMOP) of femur or skull marrow in steady state and infection. Statistics by multiple unpaired t-test followed by the Holm-Šídák, n=4-6, data represents three independent experiments. ns = not significant, ***p < 0.001. **R:** Numbers of monocytes of femur or skull marrow in steady state and infection Statistics by multiple unpaired t-test followed by the Holm-Šídák, n=4-6, data represents at least two independent experiments. ns = not significant, *p < 0.05.

These findings indicate that, unlike classical inflammatory monocyte trafficking, homeostatic monocyte access to the dura occurs largely through a CCR2-independent mechanism, whereas CCR2 becomes critical during infection-induced monocyte mobilization. The persistence of monocytes in the dura of *Ccr2^-/-^* mice consequently raised the question of their anatomical origin.

Previous reports have highlighted the skull marrow as a primary source for monocytes and B-cells replenishing the dura in homeostasis (*12*, *33*). In order to revisit this finding and evaluate the contribution of the adjacent skull marrow during GBS infection, we transplanted bone marrow from either *Cx3cr1^+/gfp^* or *β-Actin-GFP* mice, and in both cases, monocytes and monocyte-derived DM coming from the donor were GFP-labeled. Recipient mice were subjected to either full body irradiation or head-shielded irradiation, enabling us to discriminate between local (skull) versus peripheral (femur) bone marrow contribution to the dural monocyte pool (Fig. 3C). In fully irradiated mice, Ly6C^hi^ monocyte populations reached close to 100% chimerism after 8 weeks post transplantation in all assessed compartments (Fig. 3D, S3D). In contrast, femurs of head-shielded mice showed an incomplete chimerism as previously reported (*34*), while the skull marrow, which was predominantly protected by head-shielding, had significantly less contribution of donor-derived cells (Fig. 3D, S3D).

To understand donor-cell integration dynamics in dura and the bone marrow sites, we calculated relative chimerism rates by comparing the ratios of GFP-positive monocytes from dura to skull bone marrow or femur bone marrow (Fig. 3E). In fully irradiated mice, there was no difference in monocyte populations at the three sites (Fig. 3F, S3E, S3F). In head-shielded mice, relative chimerism showed that the dural monocytes closely mirrored the skull marrow chimerism (Fig. 3F). In contrast, femur monocytes had almost two times higher engraftment compared to dural monocytes (Fig. 3F). Considering the relatively short lifespan of Ly6C^hi^ monocytes (*35*), these results strongly suggested that, in steady state, the skull marrow dominated over peripheral bone marrow as monocyte reservoir for the dura. Moreover, whereas more than 75% of DM were replaced by donor-derived cells following full body irradiation, we found only 5% donor-derived DM 8 weeks post irradiation with head-shielding (Fig. 3G). For comparison, fate-mapping experiments indicate that ∼24% of DM are normally replaced by monocytes over this timeframe (Fig. S2A), suggesting that DM replacement is largely supported by skull-derived monocytes in this setting.

To determine whether the skull bone marrow supply of monocytes persists during infection, we infected head shielded chimeras 8 weeks after bone marrow transplantation with GBS (Fig. 3H). The chimerism of the skull and femur monocytes were similar in infected and non-infected mice (Fig 3K, S3H). In contrast, the percentage of donor-derived Ly6C^hi^ monocytes in the dura significantly increased upon infection, indicating an accelerated recruitment from other (non-skull) bone marrow sources (Fig. 3I, 3J, 3K, S3H, S3I). While monocytes were not detectable in the control brain, infiltrating monocytes showed a comparable chimerism to monocytes in the dura and femur upon infection in head-shielded mice, suggesting the peripheral influx in both compartments (Fig. S3H). Therefore, the accelerated replacement dynamics under infection indicated an extensive recruitment of monocytes from peripheral sources, without a preferential anatomical bone marrow niche (Fig. 3I). Additionally, the acceleration of DM replacement by donor-derived monocyte progenitors pointed to a faster monocyte-to-Mac differentiation in the acute infection (Fig. 3L). Together, these results suggest that the previously proposed CCR2-independent homeostatic replenishment of dura monocytes from the skull marrow does not suffice to meet the highly increased demand for monocytes in streptococcal meningoencephalitis. In addition and as reported before (*34*), irradiation let to a minor engraftment of donor-derived cells to the brain MG pool, which was highly reduced in head shielded chimera (Fig. S3G, S3J). Donor-derived cells in the infected head-shielded chimeras were mostly localized in the leptomeninges and were not observed integrating into the MG pool (Fig. S3J).

Given these dynamic shifts in monocyte sourcing, we next investigated whether GBS infection alters myelopoiesis within the skull and femoral bone marrow compartments. Previous studies have demonstrated that bacterial infections can skew haematopoiesis toward emergency myelopoiesis (*36*). In our model, GBS was detected in both skull and femur bone marrow, suggesting direct bacterial engagement with hematopoietic niches (Fig. 3M, S3K). While the numbers of common myeloid progenitors (CMP) remained unchanged between control and infected mice (Fig. 3N), granulocyte-monocyte progenitors (GMP) were significantly depleted in both bone marrow compartments during infection (Fig. 3O, S4A). Furthermore, monocyte progenitors and monocyte-dendritic progenitors (MDP) were significantly reduced in the skull bone marrow, whereas they were only slightly reduced in the femur (Fig. 3P, Q, S4B). Despite these changes at the progenitor level, monocyte numbers in both skull and femoral bone marrow remained unchanged during infection (Fig. 3R).

These findings indicate that GBS infection imposes distinct pressures on myeloid populations within different bone marrow reservoirs. While both sites participate in emergency myelopoiesis, the skull marrow appears particularly vulnerable to progenitor depletion, potentially reflecting its rapid mobilization of monocytes to the connected dura.

### 3.4 GBS infection reprograms monocyte ontogeny toward MDP-derived lineages

Having observed a site-specific depletion of myeloid progenitor subsets prompted us to investigate whether GBS infection alters the ontogeny of monocytes originating from these sites. Under steady-state conditions, most monocytes arise from GMP derived from CMP, although a smaller subset can develop from MDP (*37–39*)(Fig. 4A). To distinguish between these developmental pathways, we utilized *Ms4a3^Cre^;R26^Tomato^* reporter mice, in which GMP-derived monocytes (GMP-Mo) and neutrophils are irreversibly labeled (*38*), whereas MDP-derived monocytes (MDP-Mo) remain Tomato-negative. During GBS infection, MDP-Mo frequencies increased in both skull and femur bone marrow (Fig. 5B), however, this shift was significantly more pronounced in the skull, consistent with the site-specific progenitor depletion due to emergency myelopoiesis, as described above (Fig. 3O-R). Importantly, neutrophils (Fig. S5A) and GMP-derived monocyte progenitors, such as common monocyte progenitor (cMoP), remained labeled in both skulls and femurs during homeostasis and infection, confirming that altered Tomato labeling reflected genuine changes in monocyte ontogeny rather than infection-induced recombination artifacts (Fig. S5C). On the other hand, MDPs remained Tomato-negative (Fig. S5B, S5F). Furthermore, to independently verify monocyte labeling, we used *Cxcr4^CreERT2^;R26^Tomato^* mice, in which TAM induction labels all HSC-derived cells (*28*) (Fig. 4A). Indeed, there was no difference in Tomato-labeling of monocytes in skull, femur or dura in infection (Fig. S5D, E).

**Fig. 4:**
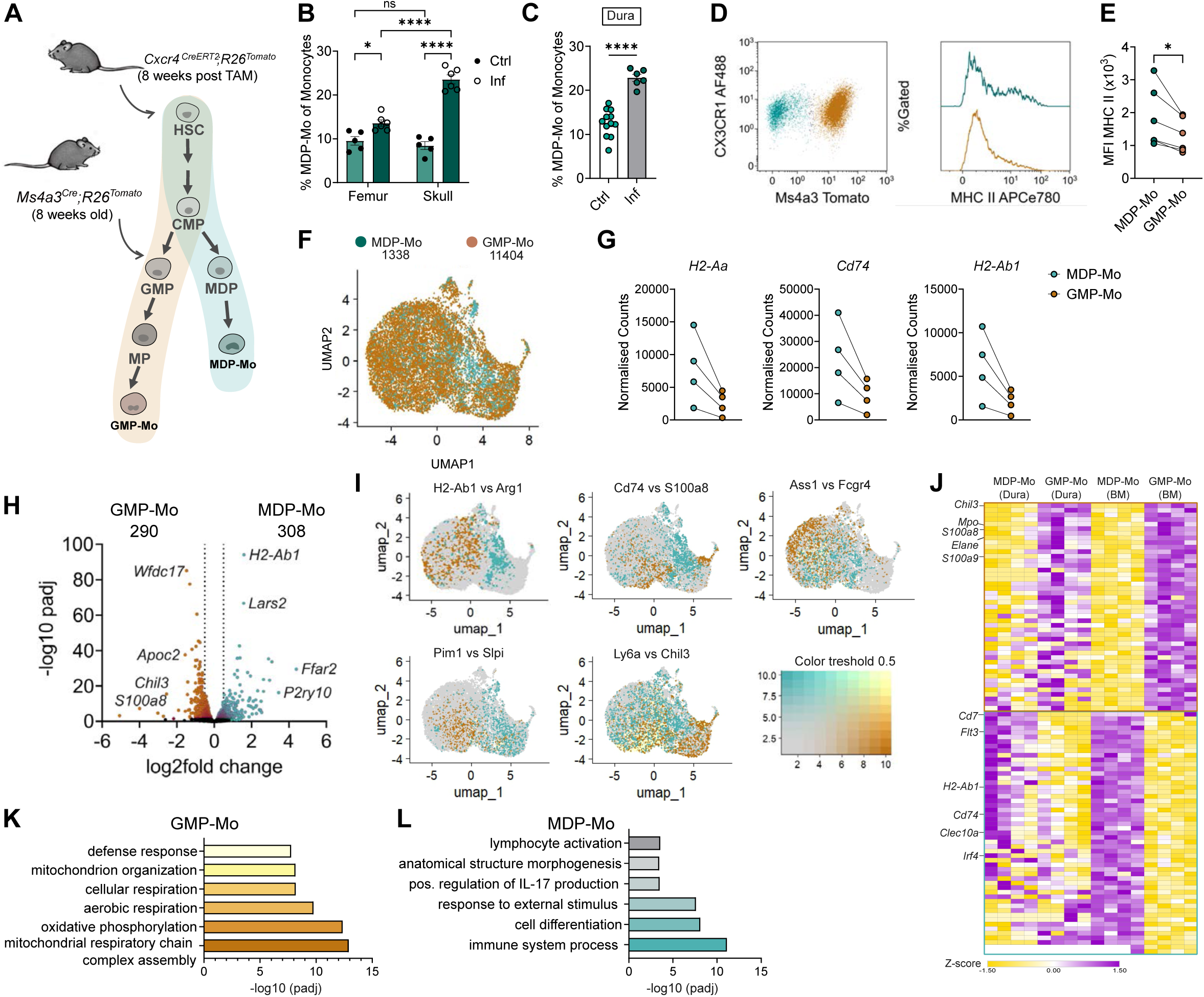
GBS infection reprograms monocyte ontogeny toward MDP-derived lineages. **A:** Scheme of either GMP or MDP-derived monocyte lineages and respective mouse models used to either label GMP-derived cells (*Ms4a3^Cre^;R26^Tomato^*) or all hematopoietic cells _(*Cxcr4*_*_CreERT2;R26Tomato_*_)._ **B**: Percentage of MDP-Mo in skull and femur bone marrow from 8-week old *Ms4a3^Cre^;R26^Tomato^*mice. n=5-6, from two independent experiments. Statistics by two-way ANOVA and Tukey’s multiple comparisons test. ns=not significant, *p<0.05,****p<0.0001. **C**: Percentage of MDP-Mo from dura of *Ms4a3^Cre/+^;R26^Tomato/+^* mice. Statistics by unpaired t-test, n=6-12, from three independent experiments. ****p<0.0001. **D:** Representative flow cytometry plots showing histograms of MHC-II expression in MDP-Mo (blue) and GMP-Mo (orange). **E:** The difference in MHC-II expression in MDP-Mo and GMP-Mo, measured as MFI from figure D. Statistics by paired t-test, n=6, from two independent experiments. **F:** UMAP representation based on gene expression profiling of sorted GMP-Mo and MDP-Mo from infected dura of *Ms4a3^Cre^;R26^Tomato^*mice. The number of analyzed cells per condition is indicated. **G:** Normalized counts of *H2-Aa*, *Cd74* and *H2-Ab1* in MDP-Mo and GMP-Mo, from pseudo-bulk analysis. **H:** Volcano plot from pseudo-bulk analysis comparing DEG in MDP-Mo (right) and GMP-Mo (left). DEG are defined as padj<0.05 and log_2_(fold change)>0.5. The number of DEGs is indicated. **I:** Feature maps showing selected genes differentially upregulated in GMP-Mo (orange) or MDP-Mo (blue), while cells expressing both are marked in yellow. **J:** Heatmap comparing DEG from GSE262369 (MDP-Mo and GMP-Mo from steady state bone marrow) with pseudo-bulk analysis from infected dura MDP-Mo and GMP-Mo. **L-M:** GO Terms for biological processes in GMP-Mo **(L)** or MDP-Mo **(M**), with exclusions of the genes overlapping with GSE262369. Only some GO Terms are represented. The whole list can be found in Suppl. Table 1.

Next, we analyzed if this ontogenic skewing was also evident in the dura during infection. Indeed, dural monocytes showed a marked increase in Tomato-negative MDP-Mo in *Ms4a3^Cre^;R26^Tomato^* mice (Fig. 4C). Of note, these MDP-Mo expressed CX3CR1 and displayed higher MHC-II levels compared to GMP-Mo (Fig. 4D, E).

To further delineate differences of MDP-Mo and GMP-Mo in the infected dura, we performed single cell RNA sequencing. Monocytes were gated as Lin^neg^CD45^+^CD11b^+^Ly6C^hi^CX3CR1^+^, with Tomato used to distinguish between lineage origins (Fig. S5G). UMAP embedding did not separate the two lineages into discrete clusters, indicating substantial transcriptional convergence in the inflamed dural environment (Fig. 4F). Nonetheless, pseudo-bulk differential expression analysis revealed clear lineage-associated signatures, including higher MHC-II gene expression in MDP-Mo, consistent with flow cytometry (Fig. 4G). In total, 308 genes were differentially expressed in MDP-Mo and 290 in GMP-Mo (Fig. 4H).

Comparison with a DEGs from published dataset of bulk RNA-seq from steady-state bone marrow–derived MDP-Mo and GMP-Mo (GSE262369) (*39*) revealed partial overlap in signature (Fig. 4I, 4J, S5H, I, J).

We then focused on DEGs that did not overlap with the steady-state dataset, reasoning that these might represent infection-associated transcriptional programs. Gene ontology analysis showed that GMP-Mo upregulated pathways related to mitochondrial respiration, oxidative phosphorylation, and antimicrobial defense, consistent with an energetically demanding inflammatory response (Fig. 4K). In contrast, MDP-Mo upregulated GO categories linked to immune system processes, cell differentiation, response to external stimuli, and positive regulation of IL-17 production (Fig. 4L).

Together, these findings indicate that GBS infection not only alters the quantity of monocytes recruited to the dura but also reshapes their developmental origins and transcriptional states, with skull bone marrow producing disproportionately more MDP-Mo under inflammatory pressure. This ontogenic reprogramming likely reflects differential stress imposed on bone marrow niches and may contribute to functional diversification of monocyte responses at the CNS interfaces during bacterial meningitis.

### 3.5 Post-infection monocyte ontogeny diverges between dura and brain

The infection-induced reprogramming of monocyte ontogeny, together with the high turnover of dural macrophages, raised the question of how the CNS myeloid landscape is restored once bacterial infection is resolved. Given that MDP-Mo expanded disproportionately during acute GBS infection in dura compared to the steady-state, we hypothesized that monocyte dynamics and their contribution to tissue macrophage pools might substantially change during the recovery phase. GBS meningitis is fatal in mice without treatment, mirroring the clinical situation in neonates. To model successful therapeutic intervention, we treated infected mice with ampicillin and azithromycin for two weeks, beginning at 2 dpi (Fig. 5A). Bacteria became undetectable in blood, brain and liver already at 7 dpi, confirming complete resolution of infection.

**Fig. 5:**
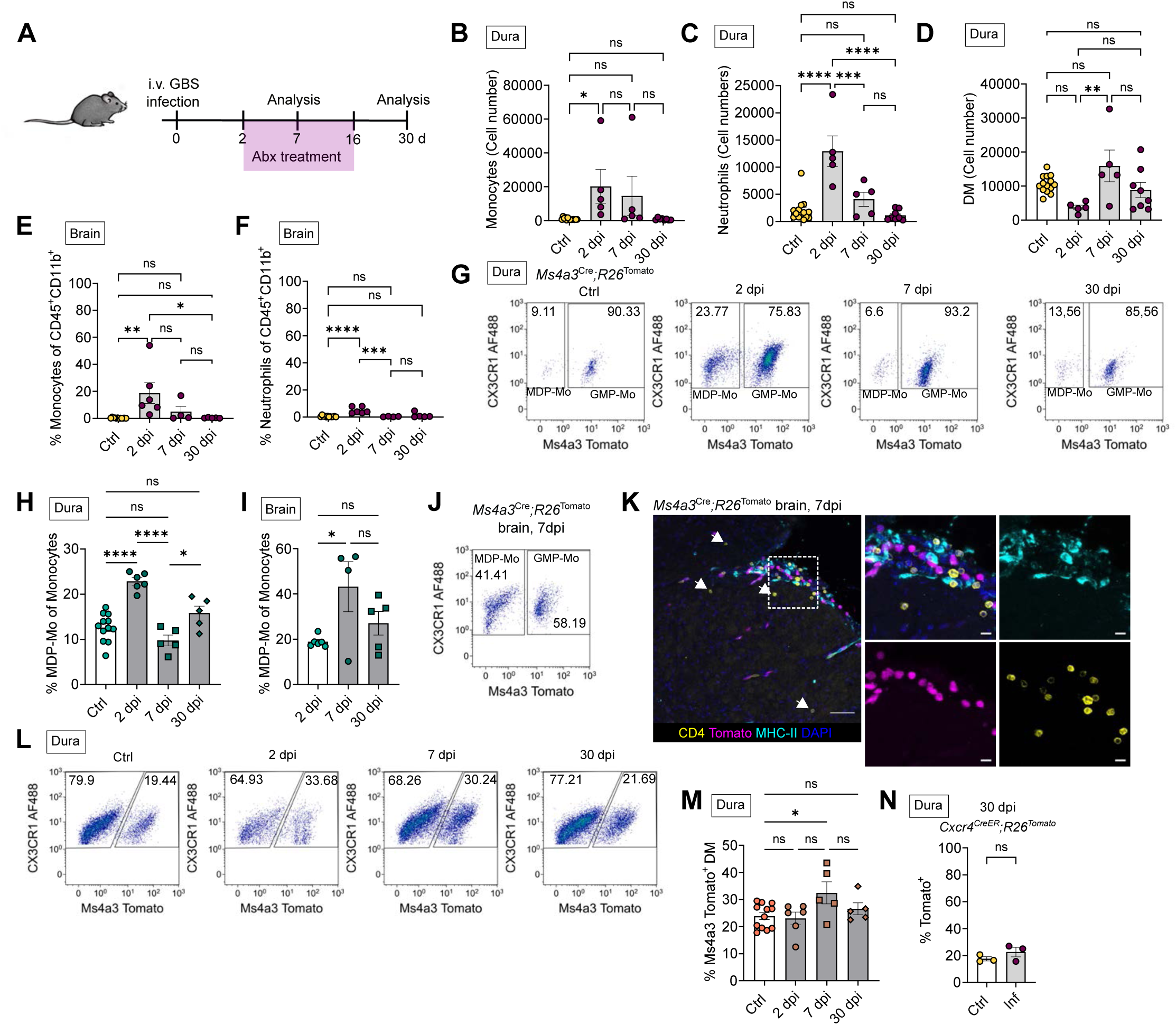
Post-infection monocyte ontogeny diverges between dura and brain. **A:** Scheme of experimental setup for antibiotic treatment. Control mice were not infected, but were treated with antibiotics for the indicated time. **B-D**: Monocyte (B), neutrophil (C) and DM (D) numbers in dura at 2, 7 and 30 dpi. Statistics by one-way ANOVA, followed by Tukey’s multiple comparisons test. N=5-14, from at least three independent experiments. **E-F:** Monocyte (E) and neutrophil (F) frequences in the brain at 2, 7 and 30 dpi. Statistics by one-way ANOVA, followed by Tukey’s multiple comparisons test. n=4-12, from at least two independent experiments. **G:** Flow cytometry plots representing changes in MDP-Mo and GMP-Mo frequencies in dura over time, from *Ms4a3^Cre^;R26^Tomato^* mice. Percentages of MDP-Mo and GMP-Mo are written in the gates. **H**: Percentages of MDP-Mo out of all monocytes in dura. Statistics by one-way ANOVA, followed by Tukey’s multiple comparisons test. n=5-12, from at least three independent experiments. **I:** Percentages of MDP-Mo out of all monocytes in brain. Statistics by one-way ANOVA, followed by Tukey’s multiple comparisons test. n=4-6, from at least two independent experiments. **J:** Representative flow cytometry plot showing MDP-Mo and GMP-Mo frequencies in brain at 7 dpi, from *Ms4a3^Cre^;R26^Tomato^*mice. Percentages of MDP-Mo and GMP-Mo are written in the gates. **K:** Representative microscopy image of sagittal brain section from *Ms4a3^Cre^;R26^Tomato^*mice at 7 dpi. Dasher rectangle represents the magnified picture on the right. Arrows point to the parenchymal CD4 T-cells. Scale bar (left)= 50 µm. Scale bar (right)= 10 µm. **L:** Flow cytometry plots representing changes in Tomato-positive and negative DM in dura over time, from *Ms4a3^Cre^;R26^Tomato^*mice. **M:** Percentages of Tomato-positive DM in dura over time, from *Ms4a3^Cre^;R26^Tomato^*mice. Statistics by one-way ANOVA, followed by Tukey’s multiple comparisons test. n=5-12, from at least two independent experiments. **N:** Percentages of Tomato-positive DM in dura at 30 dpi, from *Cxcr4^CreERT2^;R26^Tomato^* mice. Statistics by Student t-test, n=3.

By 7 dpi, monocyte and neutrophil numbers in the dura were only modestly elevated and had fully returned to baseline by 30 dpi (Fig. 5B, C). In contrast, dural macrophages expanded significantly between 2 and 7 dpi (Fig. 5D), indicating a regenerative wave of macrophage replenishment during recovery. In the brain, monocytes and neutrophils normalized by 7 dpi (Fig. 5E, 5F), although at that timepoint, monocytes persisted at low levels in the leptomeninges and parenchyma (Fig. 5E, K), suggesting ongoing immune activity beyond bacterial clearance.

Strikingly, lineage analysis revealed divergent kinetics of MDP-derived monocytes in the dura versus the brain. In the dura, MDP-Mo peaked at 2 dpi and rapidly declined to homeostatic levels during clearance (Fig. 5G, H). In contrast, brain MDP-Mo accumulated progressively and peaked at 7 dpi, constituting over 40% of the monocyte pool at this time (Fig. 5I, J). Because MDP-Mo expressed higher MHC-II than GMP-Mo and were enriched for GO terms associated with lymphocyte activation and IL-17 production, we hypothesized that their delayed accumulation in the brain may reflect a role in antigen presentation and T-cell activation. Indeed, CD45^+^CD11b^neg^ lymphocytes increased specifically in the brain (Fig. S6A), but not the dura (Fig. S6B), at 7 dpi. Histological analysis of brain 7 dpi showed accumulation of CD4^+^ T-cells in proximity of MHC-II^+^ cells, of which most are found in CNS-border areas such as the ventricles or the leptomeninges, but were also present in the brain parenchyma (Fig. 5K, S6D). On the other hand, at 2 dpi, accumulation of monocytes was found in leptomeninges with very rare CD4^+^ T-cell incidence, while at 30 dpi neither monocytes nor CD4^+^ T-cells are present in the brain (Fig. S6C).

We next asked whether GMP-derived monocytes contribute to restoration of the DM niche. Tomato^+^ DM in *Ms4a3^Cre^;R26^Tomato^*mice transiently increased in the dura at 7 dpi but were back to steady state levels at 30 dpi, indicating they only contribute to the DM compartment transiently (Fig. 5L, M). Since MDP-Mo and MDP-derived Macs cannot be fate-mapped with currently available tools, we assessed long-term DM maintenance using *Cxcr4^CreERT2^;R26^Tomato^*mice induced one week before infection. Tomato^+^ DM frequencies were indistinguishable between infected and control mice at 30 dpi (Fig. 5N), demonstrating that the DM niche is ultimately restored by long-lived tissue-resident macrophages rather than recruited monocyte progeny. Similarly, microglia remained Tomato-negative, indicating that CNS parenchymal macrophages do not undergo substantial monocyte-derived replacement during or after GBS infection (Fig. S6E).

Thus, while the dura and brain recruit distinct monocyte populations during infection and early recovery, both compartments ultimately revert to a resident cell–dominated homeostasis.

## 4. DISCUSSION

Acute streptococcal meningoencephalitis warrants major prevention efforts worldwide, due to its substantial consequences for cognition and motor function in surviving patients (*40*). The uniqueness of the disease in an individual’s life is reflected by the long-standing concept of a largely immune-privileged brain, which - unlike other internal organs, such as the liver and kidneys, does not come into incidental contact with pathogens and peripheral immune cells (*41*). Accordingly, most mechanistic insights have focused on MG, the resident immune cells of the CNS, largely because of the assumption that they are the primary mediators of immune defense within the CNS interior (*16*, *18*, *42*, *43*).

In contrast, very little information is available on the role of CAM in this context, despite the fact that clinically, CNS borders play a central role in disease manifestation. This discrepancy is probably due to the fact that until recently it was not possible to reliably distinguish between CAM and parenchymal MG, and owed to the paucity of robust rodent meningitis models in which bacteria are not injected directly into the CNS.

Lately, there has been increasing evidence that macrophages residing in brain barriers, particularly in the meninges, play an important role in various neuropathologies (*4*, *5*, *44*). In addition, the recent discovery of direct communication between the dura mater and skull bone marrow raises the question of the role of distinct bone marrow sites in disease (*11–13*, *45*, *46*).

Accordingly, we have combined the latest tools in myeloid cell fate-mapping and multidimensional analysis with a mouse model in which streptococci circulate systemically before entering the CNS, as in a natural disease. In this model, we demonstrate that bacteria accumulate primarily in the leptomeninges and dura, with minimal penetration into the brain parenchyma, which aligns with emerging evidence that the meninges behave as a highly responsive immunological interface (*47*, *48*).

Unlike perivascular and leptomeningeal macrophages, DM are replenished to a higher percentage by monocytes in steady state, reaching around 45% DM replacement after 4 months. These dynamics changed rapidly during the infection, characterized by the disappearance of resident DM and a faster differentiation of incoming monocytes into a Ly6C^+^CD206^+^MHC-II^+^ macrophage population, which is not discernible in steady state. scRNA-seq data from infected mice confirmed the emergence of infection-associated monocytes which upregulated *Mrc1* and MHC-II genes proportionally to the bacterial burden found in the brain. Additionally, adoptive transfer of monocyte progenitors validated the capacity of monocytes to rapidly assume macrophage features in the infected dura.

On the other hand, bacterial burden positively correlated to the loss of different DM clusters. It is conceivable that these resident DM populations are the first responders in the early phase of infection, leading to cellular stress and ultimately, cell death. This observation is consistent with the so-called “macrophage disappearance reaction” which has been described in inflammation of the peritoneal cavity (*49*, *50*), the liver (*31*) and other tissues (*51*) and which might serve to initiate infection control. Infection-induced cell death of resident Macs has also been found in the meninges in viral infection (*4*), accompanied by the engraftment of monocytes. Consistent with this, the upregulation of canonical necroptosis genes *Mlkl, Ripk1* and *Zbp1* in DM clusters in infection points to a likely mechanism of DM loss in our model.

The skull bone marrow can serve as a reservoir for myeloid cells, which directly supplies the dura with monocytes (*12*). This finding is particularly interesting considering the great demand for monocytes in meningoencephalitis, not only for the induction of inflammation, but also to serve as potential Mac precursors in a niche emptying due to Mac death.

By using head-shielded bone marrow chimeras, we confirm the recent finding that the skull marrow supplies the adjacent dura with monocytes in steady state (*12*). Remarkably, head-shielded mice had less than 8% of donor-originating DM, whereas our fate-mapping experiments revealed that monocyte replacement would reach around 24% over 8-week period. These results point to the skull marrow as the primary source of monocytes for DM renewal in homeostasis. However, infection alters homeostatic pattern of monocyte egress and differentiation. In infected head-shielded mice, chimerism rates were significantly higher in dura and brain, suggesting that skull marrow could not adequately meet the high demand for monocytes in meningoencephalitis and that other sources of hematopoietic progenitor cells were required. This observed „recruitment threshold“ during infection was paralleled by changes in skull marrow composition, particularly in monocytes and monocyte precursors, including GMP-derived monocyte progenitors (MP) and MDP. Together with the observed presence of GBS in the skull marrow, this implies the initiation of emergency hematopoietic programs. Indeed, our data support this shift in myelopoiesis. While the homeostatic dura hosts classical and non-classical monocytes, we found that in infection monocyte numbers substantially expanded, with a significant influx of transcriptionally distinct infection-associated monocytes. Furthermore, we show that, while the steady state dura can be replenished by monocytes in CCR2-independent manner, CCR2 was required for the infection-associated monocyte influx into the dura. It seems noteworthy that we did not find evidence that these infection-associated monocytes contributed to bacterial trafficking, as *Ccr2^-/-^* mice did not have a difference in bacterial load compared to *Wt* animals. This is consistent with studies that have identified numerous genes in GBS which facilitate bacterial adhesion to the endothelium, thereby enabling transcytosis and potentially crossing of CNS barriers (*21*, *52*, *53*). Our lineage tracing using *Ms4a3^Cre^* mice uncovered infection-driven changes in monocyte ontogeny. While GMP-derived monocytes dominate in steady state, infection induced a substantial increase in MDP-derived monocytes, especially in skull marrow and the dura. MDP-Mo displayed elevated MHC-II and transcriptional programs associated with lymphocyte activation, differentiation, and IL-17 pathways. Whereas GMP-Mo expressed metabolic and antimicrobial pathways consistent with classical inflammatory responses, MDP-Mo appeared positioned to modulate adaptive immunity. Indeed, after the clearance of the infection by using antibiotics, we found more than 40% of MDP-Mo in the brain and leptomeninges, as well as CD4+ T-cells. Nonetheless, neither MDP-Mo nor GMP-Mo contributed lastingly to the DM or MG pools during recovery. Indeed, our *Cxcr4^CreERT2^* fate-mapping analysis indicates that surviving tissue-resident cells restored the DM niche in the long term.

Nevertheless, additional work is needed to understand whether changes in monocyte ontogeny modify disease progression and to determine the cues that govern the transition from GMP- to MDP-derived monocyte production. Furthermore, GBS meningoencephalitis predominantly affects neonates and young infants, whose barrier tissues and immune system undergo rapid changes and differ from adults. These developmental differences likely influence the magnitude, kinetics, and cellular composition of the meningeal immune response. Thus, our model in 6–12 week old mice may not fully reflect the unique vulnerabilities of the perinatal CNS relevant to neonatal GBS infection. Future studies applying this infection paradigm to neonatal or early postnatal mice will be important to determine how age-dependent differences in barrier integrity, macrophage maturation, and monocyte ontogeny shape the trajectory of meningeal infection.

In summary, we have integrated fate mapping, microscopy and transcriptomic analysis to highlight a multilayered immune architecture at the CNS border during streptococcal meningoencephalitis. These insights reshape our understanding of myeloid dynamics during bacterial CNS infection, suggesting that monocyte origin, differentiation trajectory, and anatomical context collectively determine their function in disease. Furthermore, the vulnerability of skull marrow progenitors to bacterial invasion and depletion suggests that local hematopoietic niches may influence susceptibility or recovery potential in CNS infections.

## 5. MATERIALS AND METHODS

### 1. Study design

This study aimed to decipher changes in the myeloid populations in the CNS during streptococcal meningoencephalitis, specifically focusing of microglia, dural macrophages and monocytes. For that purpose, we used a combination of different fate mapping mouse models, as well as microscopy, flow cytometry and transcriptomic analysis. Group sizes and number of repetitions are indicated in the figure legends.

### 2. Mice

Both female and male mice from C57BL/6 genetic background were used at 6 to 12 weeks of age. C57BL/6J and C57BL/6N mice were purchased from Jackson Laboratories (USA) or Charles River Laboratories (Germany) and bred in the local animal facility. CD45.1-mice (B6.SJL-Ptprca Pepcb/BoyJ) were purchased from Jackson Laboratories. *Cx3cr1^gfp/+^* mice were obtained as a kind gift from Steffen Jung (Weizmann Institute, Israel). *β-Actin gfp+/− (C57BL/6-Tg(CAG-EGFP)131Osb/LeySopJ)* and *Ms4a3^Cre^* (C57BL/6J-Ms4a3em2(cre)Fgnx/J) mice were purchased from Jackson Laboratories. *Mrc1^CreERT2^* mice were provided by Marco Prinz (Institute of Neuropathology, Freiburg) as a kind gift. *Cxcr4^CreERT2^* mice were provided by Ralf Stumm (Neuropharmacology, Jena) as a kind gift. *Ccr2^-/-^* (B6.129S4-Ccr2tm1Ifc/J) mice were bought from Jackson Laboratories (USA).

Mice were bred in animal facilities of the University of Freiburg under specific pathogen-free conditions and were kept in groups of a maximum of five animals per cage at defined environmental conditions. Food and water were available *ad libitum*, and day/night cycles were set to 12 h. All animal experiments were approved by the Federal Ministry for Nature, Environment and Consumer’s protection of the state of Baden-Wuerttemberg (proposal numbers G-19/031 and G23/008).

### 3. Mouse interventions

Induction of Cre recombinase by tamoxifen has been explained previously(*30*). Briefly, mice were anesthetized by isoflurane inhalation. 4mg TAM in 200μl corn oil (both Sigma) was injected subcutaneously in *Cxcr4^CreERT2^* or *Mrc1^CreERT2^* mice at two time points 48h apart. As control, littermates with respective floxed alleles, but without Cre recombinase expression, were treated in the same fashion.

For bone marrow transplantations, recipient mice were anesthetized using ketamine (100 mg/kg body weight) and xylazine (10 mg/kg body weight) administered intraperitoneally. Mice were then head-shielded with a 6 mm lead plate, or not shielded and lethally irradiated with 9 Gy. Subsequently, mice received 5x10^6^ donor-derived bone marrow cells intravenously. Mice were analyzed 8 weeks post transplantation.

Mice treated with antibiotics were injected i.p. with ampicillin (100 μg/g body weight) and azithromycin (50 μg/g body weight) dissolved in 100 μl sterile water at 2dpi, after which they continued to receive antibiotics via the drinking water for up to two weeks (0.4 g/L ampicillin and 0.2 g/L azithromycin). Control mice were injected with PBS, then treated with antibiotics. Mice were analyzed either at 7 dpi or 30 dpi.

### 4. GBS culturing and infection

Prior to infection, GBS strain BM110 was streaked on a blood agar plate (Columbia agar + 5% sheep blood; BioMérieux) and incubated at 37°C overnight. One colony from the blood agar plate was transferred into 5 ml of Todd-Hewitt broth and incubated at 37°C overnight. The overnight culture was inoculated into 200 ml warm Todd-Hewitt broth. Bacteria were grown to the mid-log phase, and bacterial concentration was determined with an optical spectrophotometer (OD600 = 0.4) and confirmed by serial dilutions plating onto a blood agar plate. Afterwards, bacteria were washed twice with PBS and resuspended in PBS at 10^9^ CFU/ml. A total of 50 µl (5 x 10^7^ CFU) suspension of bacteria or vehicle (PBS) was injected into the retro-orbital venous plexus. For this procedure, mice were anesthetized by isoflurane inhalation.

### 5. CFU assay

Mice were sacrificed by CO_2_ inhalation after which the blood was drawn from the heart. Mice were then transcardially perfused with ice-cold sterile PBS. Liver, spleen and brain were collected into sterile plastic bags, and mechanically dissociated. Femur was flushed through a 70 µm cell strainer using 2.5-3 ml sterile PBS in the syringe, and 20 µl of suspension was plated on Granada agar (BD). Skull marrow was extracted as previously described(*12*), with small modifications. After the removal of dura, the calvarium was cut into small pieces into 1 ml sterile PBS using sterile scissors and plated on Granada agar. Dura was incubated for 20 min at 37°C with shaking in 1 ml of sterile RPMI containing 2.5 mg of Collagenase IV after which 20 µl of suspension or serial dilution was plated on the Granada agar. To control sterility of the CFU assay, tissues of the control animals were plated in parallel, with no bacterial growth from any analyzed organ.

### 6. Cell extraction

Blood was drawn from the heart. Erythrocytes were lysed in RBC Lysis buffer solution (eBioscience™ 1X RBC Lysis Buffer). The remaining cells were washed in FACS buffer (PBS + 1% FBS + 2 mM EDTA).

Dura mater was peeled using forceps from calvaria and placed into a digestion solution (2.5 mg/ml collagenase IV in RPMI) and incubated for 20 min at 37°C shaking, after which the solution was filtered through 70 µm strainer (BD) and washed with FACS buffer.

The remaining calvaria was cleaned from leftover muscle and fascia, placed into a 2 ml Eppendorf tube with 1 ml PBS and diced with scissors. The bone marrow was filtered through a 70 µm strainer and washed with FACS buffer.

Femur was cleaned from the leftover muscle, and the bone marrow was flushed with PBS using a 26G needle through a 70 µm strainer.

Brain was homogenized and filtered through a 100 µm strainer. Myelin was removed using 37% Percoll gradient and centrifugation for 30min at 300g RT, with no acceleration and brake.

Samples were washed and stained with respective antibodies. FcγII/III receptors were blocked by prior 10 min incubation with Fc block (anti mouse CD16/32 antibody Clone 93, Biolegend) on ice.

Samples used for bulk RNA-seq (dura and brain) were processed in the same way to avoid potential bias of Mac activation due to enzymatic digestion. Shortly, brain and dura were incubated for 20 min at 37°C shanking in 2.5 mg/ml collagenase IV, after which they were filtered through 70 µm strainer and washed with PBS. The pellet was resuspended using 37% Percoll and centrifuged for 30 min at 300g RT, without the acceleration and brake.

### 7. Flow cytometry and FACS

Cells were stained with the indicated antibodies according to manufacturer’s instructions. Flow cytometry was performed on a 10-color flow cytometer Gallios™ (Beckman Coulter) or Cytoflex S™ (Beckman Coulter) and FACS was done on Moflo^TM^ (Beckmann Coulter) or CytoFlex™ (Beckman Coulter).

The following anti-mouse antibodies were used for surface staining:

anti-mouse CD3e APC Clone 145-2C11 (BioLegend)

anti-mouse CD16/32 PE Cy7 Clone 93 (eBioscience)

anti-mouse CD19 APC Clone 1D3/CD19 (BioLegend)

anti-mouse CD34 PE Clone HM34 (BioLegend)

anti-mouse CD45 eFluor450 Clone 30-F11 (eBioscience),

anti-mouse CD45 FITC Clone 30-F11 (eBioscience)

anti-mouse CD45 PerCP Cy5.5 Clone 30-F11 (eBioscience)

anti-mouse CD45.1 FITC Clone A20 (eBioscience)

anti mouse CD45.2 PerCP Cy5.5 Clone 104 (eBioscience)

anti mouse CD11b FITC Clone REA592 (Miltenyi)

anti-mouse CD11b PE-Cy7 Clone M1/70 (eBioscience)

anti-mouse CD11b APC-eFlour780 Clone M1/70 (eBioscience)

anti-mouse CD117 BV421 Clone 2B8 (BioLegend)

anti-mouse CD117 PE Cy7 Clone 2B8 (eBioscience)

anti-mouse CD115 APC Clone AFS98 (Miltenyi)

anti-mouse CD115 BV421 Clone AFS98 (BioLegend)

anti-mouse CD127 APC Clone A7R34 (BioLegend)

anti-mouse CD206 APC Clone C068C2 (BioLegend)

anti-mouse CD206 PE Clone C068C2 (BioLegend)

anti-mouse CD206 AF488 Clone C068C2 (BioLegend)

anti-mouse CD206 PE Clone C068C2 (BioLegend)

anti-mouse CX3CR1 AF488 Clone SA011F11 (BioLegend)

anti-mouse Ly6G FITC Clone 1A8 (BD Pharmingen)

anti-mouse Ly6G APC Clone 1A8 (BD Pharmingen)

anti-mouse Ly6C BV605 Clone HK1.4 (BioLegend)

anti-mouse MHC II APC eFlour780 Clone M5/114.15.2 (eBioscience)

anti-mouse Ter119 APC Clone TER-119 (BioLegend)

anti-mouse Sca-1 AF488 Clone D7 (BioLegend)

Analysis was done using Kaluza software (v1.5a and 2.1, Beckman Coulter).

For calculating absolute numbers of cells in dura, the whole dura was acquired. For calculating absolute numbers of populations in brain, skull or femur, a defined volume of the processed tissue was acquired for a pre-defined time (30-60s) at a flow of 10 µL/min or 30 µL/min.

### 8. Adoptive transfer

Bone marrow was isolated from tibia and femur of *β-Actin-gfp^+/−^* mice, flushed with sterile ice-cold PBS and centrifuged for 6 min at 1500 rpm. Cells were resuspended in 1% BSA in PBS, and then stained and sorted as described above. Sorted monocyte progenitors were then resuspended in PBS + 2% murine serum which was obtained from the donor mice. Monocyte progenitors were injected intravenously at 1 dpi (∼2–4 × 10^5^/animal). Recipients were analyzed 1 day post transfer (2 days post infection). Blood, brain, dura and bone marrow (from femur and skull) were analyzed on a flow cytometer (Gallios™, Beckman Coulter) and processed with the Kaluza software (v2.1, Beckman Coulter) and/or by microscopy.

### 9. RNA extraction and RT-qPCR

Cells were sorted into RLT lysis buffer (Qiagen) with 1% β-mercaptoethanol. RNA was extracted with RNeasy MicroKit (Qiagen) according to the manufacturer’s instructions. RNA was reversely transcribed into cDNA using Superscript IV (Invitrogen). RT-qPCR was performed with absolute qPCR SYBR Green Mix (Thermo Fisher Scientific) using Roche Lightcycler^TM^ 175. mRNA levels were normalized to *Gapdh* as housekeeping gene.

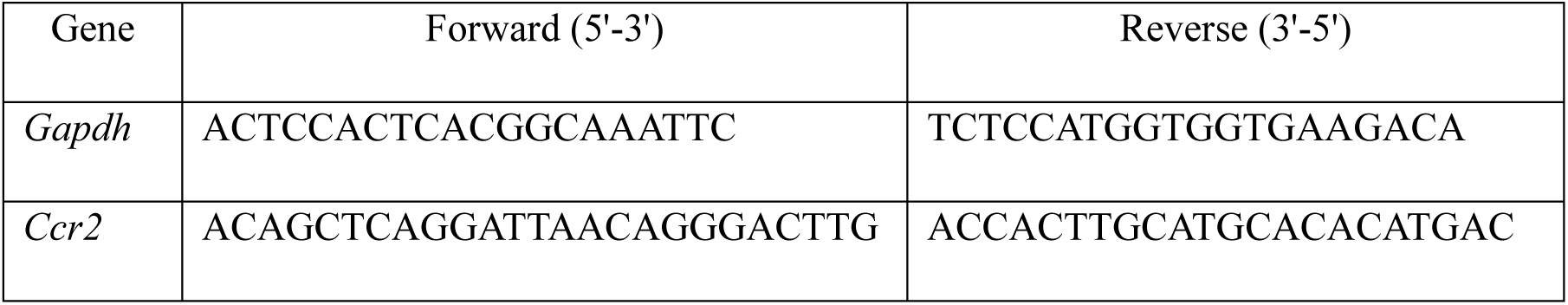

### 10. Bulk RNA sequencing

For bulk RNA sequencing, described populations were sorted into Rneasy Microkit (Qiagen) lysis buffer containing 1% β-mercaptoethanol. RNA was extracted following manufacturer’s protocol. To insure appropriate RNA amounts, samples from two mice were pooled for n=1 in both conditions. Bulk RNA sequencing was performed by CeGat (Tübingen, Germany). Sequencing was performed with 2 x 100 bp read length and 50 million clusters/probe on NovaSeq 6000 (Illumina). Raw data was provided by CeGat. Group comparison was done using DESeq2 in R. PCA plots were done using DESeq2 in R 4.3.3 and RStudio version 2023.12.1. GO terms were analyzed with the online tool “g:Profiler”(*54*). Heat maps were made using Morpheus (https://software.broadinstitute.org/morpheus).

### 11. Single-cell RNA sequencing

For scRNA Seq of myeloid population, dura mater was prepared from five infected and six control *Wt* mice as described above in section 5.

Prior FACS, cells from individual control or infected mice were labelled using the 10X 3’ CellPlex Kit Set A (PN-1000261) following manufacturer’s recommendations. CD45^+^CD11b^+^ cell population was sorted. In total, 5-6 individual animals per group were pooled after FACS. For controls, around 10 750 cells were sorted per mouse (64 500 cells in total), and for infected mice, in average 11 600 cells were sorted per mouse (46 400 cells in total). Sorting was done on separate days for control and infected mice due to technical limitations.

Sorted samples were processed using the 10X Chromium Next GEM Single Cell 3’ Kit v3.1 (PN-1000269) and 3’ Feature Barcode Kit (PN-1000262). Single cell droplets were generated using Chromium Next GEM Chip G Single Cell Kit (PN-1000127) and Chromium Controller for 3’ gene expression and cell multiplexing libraries were generated according to the manufacturer’s specifications. Library quality was assessed using Agilent TapeStation 4150 Cell-free DNA ScreenTape assay (cat. no. 5067-5630) and NEBNext Library Quant Kit for Illumina (cat. no. E7630S). The single cell libraries were sequenced using Ilumina NextSeq1000/2000, with a minimum targeted sequencing depth of 20,000 reads per cell. Fastq files were aligned using the CellRanger version 7.1.

For scRNA Seq of MDP-Mo and GMP-Mo, dura from 4 infected *Ms4a3^Cre^;R26^Tomato^* mice was processed as described above. Prior FACS, single-cell suspension of each individual mouse was labelled using the 10X 3’ CellPlex Kit Set A (PN-1000261) following manufacturer’s recommendations. Sorting of MDP-Mo and GMP-Mo was done as described in Fig. S5G. Dump gate included CD206, CD3 and CD19 (all in APC). In total, around 7500 MDP-Mo and 59400 GMP-Mo were sorted. Sorted samples were processed using the 10X Chromium Next GEM Single Cell 3’ Kit v3.1 (PN-1000269) and 3’ Feature Barcode Kit (PN-1000262). Single cell droplets were generated using Chromium Next GEM Chip G Single Cell Kit (PN-1000127) and Chromium Controller for 3’ gene expression and cell multiplexing libraries were generated according to the manufacturer’s specifications. Library quality was assessed using Agilent High Sensitivity DNA Kit (cat. no. 5067-4626). The single cell libraries were sequenced using Ilumina NovaSeq X Plus, with a minimum targeted sequencing depth of 20,000 reads per cell. Fastq files were aligned to prebuilt CellRanger reference refdata-gex-GRCm39-2024-A using the CellRanger version 9.0.1.

### 12. Single-cell RNA sequencing downstream analysis

For myeloid population scSeq from control and infected dura, downstream analyses were conducted in the R version 4.2.0 programming environment using the Seurat package version 4. Cells were retained if the mitochondrial gene content was below 5%. To exclude empty droplets or doublets we limited the number of detected genes per cell between 200 and 5,000. Total counts per cell had to exceed 500 to ensure a minimum level of transcriptional activity indicative of viable cells.

In addition, we filtered out specific classes of genes that can introduce potential biases by overrepresentation of ribosomal or pseudogenes in the scRNAseq data. Genes belonging to the murine-specific ribosomal protein large (Gm) family, and ribosomal proteins (Rpl and Rps), were excluded. These genes were identified using regular expressions to match gene names starting with "Gm", "Rpl", or "Rps" and were removed from the analysis.

Data was integrated from two separate experiments using the SCtransform algorithm within Seurat. 2000 features were identified using SelectIntegrationFeatures() and used in PrepSCTIntegration() and FindIntegrationAnchors(). These anchors were used on IntegrateData(). The top 30 PCAs were used for Umap calculation. FindNeighbors() and FindClusters() were calculated with the top 30 dimensions each. For the latter, a resolution of 0.299 was chosen as the highest resolution where every cluster was distinguishable by specific biological functions. Core markers were identified using the FindAllMarkers functions using a threshold of at least 25% of cells expressing the gene and only returning positive values. All clusters are displayed in the respective Violin plots generated by VlnPlot() and sorted according to the respective median.

For monocyte scSeq (GMP-Mo and MDP-Mo), downstream analyses were conducted in R version 4.5.1 programming environment using the Seurat package version 5.3.1. Cells were retained only if the mitochondrial gene content was below 20% and we restricted the number of detected genes per cell to be between 300 and 6,000. Furthermore, total UMI counts per cell were required to exceed 500.

In addition to cell-level filtering, we removed specific classes of genes that could introduce bias due to overrepresentation in scRNA-seq datasets. Genes corresponding to mitochondrial transcripts were identified using the regular expression pattern "(?i)^mt-", and ribosomal protein genes were detected using the pattern "(?i)^(rpl|rps)". To compare transcriptional profiles between MDP-Mo and GMP-Mo populations, pseudobulk function was used, in which counts from individual cells belonging to the same sample and lineage were aggregated to generate sample-level expression profiles. Pseudobulk count matrices were generated for each sample × lineage combination, normalized, and analyzed using DESeq2. DEG were defined as p<0.05, logFold2(Change)>0.5. For comparison with bulk RNA seq data from GSE262369, published data was analysed using DESeq2, in which DEG were defined as p<0.05, logFold2(Change)>0.5. Heat maps were made using Morpheus (https://software.broadinstitute.org/morpheus).

### 13. Immunofluorescence labelling

After transcardial perfusion with ice-cold PBS, brain was removed, fixed in 4% PFA for 24 h and cryoprotected in 30% sucrose at 4°C until it sank (1-2 days). After tissue freezing in OCT compound (Sakura, Zoeterwoude, Netherlands) on dry ice, the brain was cut in 30-40 µm sagittal cryosections. Sections were firstly permeabilised for 2h using 0.2% Triton-X in PBS, after which the samples were blocked in blocking solution containing 10% goat serum and 0.1% Triton-X in PBS. Samples were stained with mastermix containing PBS, 5% goat serum, 0.05% Triton-X and antibodies, overnight at 4°C. Dilutions of antibodies were 1:500 for Iba-1 (WACO, Japan), 1:200 for anti-GBS (Invitrogen), 1:100 for CD4 – AF488, CD74 – AF647, CD206 – PE, CD206- APC (Biolegend), 1:100 for CD45-PE (Invitrogen), 1:100 for CD31-PE (BD Pharmigen) and 1:50 for MHC-II APC (Miltenyi). After washing 5x with PBS, samples are incubated for 2h at RT with secondary antibody (goat anti-rabbit Alexa Fluor633 1:500, anti-GFP DyLight488 1:1000 for enhancing endogenous signal in *Cx3Cr1^/gfp^*mice) and DAPI (1:1000). Samples were washed 5x and mounted on SuperFrost slides using ProLong Diamond Antifade Mountant (Invitrogen). Whole mounts of dura mater were prepared as previously described(*17*), with minor modifications. After transcardial perfusion with ice-cold PBS, calvaria containing cortical dura mater was placed in 4% PFA overnight at 4°C. The dura was dissected and placed into PBS overnight at 4°C to remove any PFA. Samples were then blocked for 2h with agitation on RT using a blocking solution containing 5% goat serum and 0.1% Triton-X. Samples were stained overnight at 4°C in staining solution containing 2% goat serum, 0.1% Triton-X and primary antibodies. Primary antibodies used are as follows: anti-GBS (1:200), CD206 (1:100), CD31 (1:100). Nuclei were stained by DAPI (1:1000).

Washing was done 5x with PBS before using the secondary antibodies diluted in a staining solution. Samples were incubated for 2 h at RT. Stained samples were washed thoroughly with PBS and mounted using ProLong Diamond Antifade Mountant.

For skull and femur, the tissue was cleaned from muscle and fascia and placed in 4% PFA overnight at 4°C. Next day, the tissues were transferred into decalcification buffer (Osteosoft®, Sigma Aldrich) and shaken for 2 days in a 37°C incubator. The tissues were then washed in PBS and frozen in OCT compound. 15-20 µm cryosections were mounted on gelatin-coated slides. Cuts were rinsed in PBS and washed with PBS with 0.2% Triton X-100 for 30 min, followed by incubation in blocking and permeabilization buffer containing PBS with 0.2% Triton X-100 and 5% normal goat serum for 1 h at RT. Primary antibodies were diluted in 0.05% Triton-X and 2% NGS and incubated at 4°C overnight. The antibody dilutions were: CD45 (1:100), anti-GBS (1:200). Then, sections were washed with PBS 5x, and stained with secondary antibody and DAPI for 2 h at RT. Sections were then washed thoroughly with PBS and mounted using ProLong Diamond Antifade Mountant.

Heads were cleared from skin and mandible was removed. Heads were kept in 4% PFA at 4°C overnight. Next day, the heads were transferred into decalcification buffer and shaken for 2 days in a 37°C incubator. Heads were then coronally cut with scalpel into approximately 5 mm slices and frozen in OCT compound. 70-80 cryosections were mounted on gelatin-coated slides. The downstream processing and staining was the same as for the brain.

### 14. Confocal microscopy

Confocal microscopy was performed with LSM 880 or LSM 980 confocal microscopes equipped with a 20 × /0.8 NA Plan-Apochromat objective (Carl Zeiss Microimaging) using 1-1.5 μm optical slices. Images are displayed as maximum-intensity projections of 25-40μm thick z stacks, 1024x1024 pixel resolution.

### 15. Image analysis

Area and fluorescence intensity were measured using ZEN 3.0 (blue edition, Zeiss). For enumeration of DM per field of view, the pictures were taken on approximately same locations in each dura. CX3CR1-GFP^+^ DM were manually counted using ImageJ (version 1.54p) Cell counter.

### 16. ELISA

For brain ELISA, one brain hemisphere was weighed and stored in 1 ml of homogenization buffer (50 mM Tris pH 8, 150 mM NaCl, 5 mM EDTA) containing 10 µl phosphatase and protease inhibitors (Pierce). Brain was then homogenized using tissue lyser (Quiagen^TM^ TissueLyser) with 50/s oscillations for 5 min with 3 mm tungsten carbide beads. Tissue debris was removed by centrifugation at 20 000 g for 10 min at 4°C. Supernatants were kept at -80°C until the analysis. For plasma ELISA, blood was collected into Microvette® tube (Sarstedt) and centrifuged at 10 000 g for 10 min. Plasma was collected and kept at - 80°C until the analysis. Concentrations of IL-6, TNF, IL-1β and IFN-β were measured using mouse R&D Duo set ELISA kits according to the manufacturer’s instructions (R&D Systems).

### 17. Statistical analysis

The analysis used for each experiment can be found in figure legends, with the number of replicates and independent experiments. Differences were considered statistically significant if p-values were < 0.05. (ns, not significant; *p < 0.05; **p < 0.01; ***p < 0.001; ****p < 0.0001). Data are represented as mean ± SEM if not stated otherwise.

GraphPad prism version 10.0.0 was used for statistical analysis.

## List of supplementary materials

Supplemental figures 1-6, Supplemental Table 1

## Acknowledgments

We thank Anita Imm, Reem Alsumati, Adriana Greco and Lena Grubišić for their great technical assistance. We thank the ZTZ core facility, especially Jan Bodinek and Marie Follo for their great technical assistance. We thank the Center for Experimental Models and Transgenic Service (CEMT) of University Medical Center Freiburg for excellent technical support and assistance with the animal studies performed in this work. We thank Pedro Aniceto for his assistance with analysis in R and R Studio.

## Author contributions

Conceptualization: PH, VG Supervision: PH, DE

Formal analysis: VG, FL, LFPB, GM, S, DO

Investigation: VG, VF, AL, JN, ZMM, TM, SB, JH

Resources: PH, KPK, MP, SJ

Validation: VG, JK Visualization: VG

Writing – original draft: VG, PH

Writing – review and editing: PH, KK, JK, DE, S, FL, AL, JK, SB, VF, JN, ZMM

## Data and materials availability

Please contact the first (VG) or corresponding author (PH) if you want an access to the transcriptome data generated in this study or other information not covered in the method section.

