## Supplemental Figures for "Streptococcal meningoencephalitis dynamically shapes developmental monocyte and macrophage trajectories in the CNS"

### Fig. S1

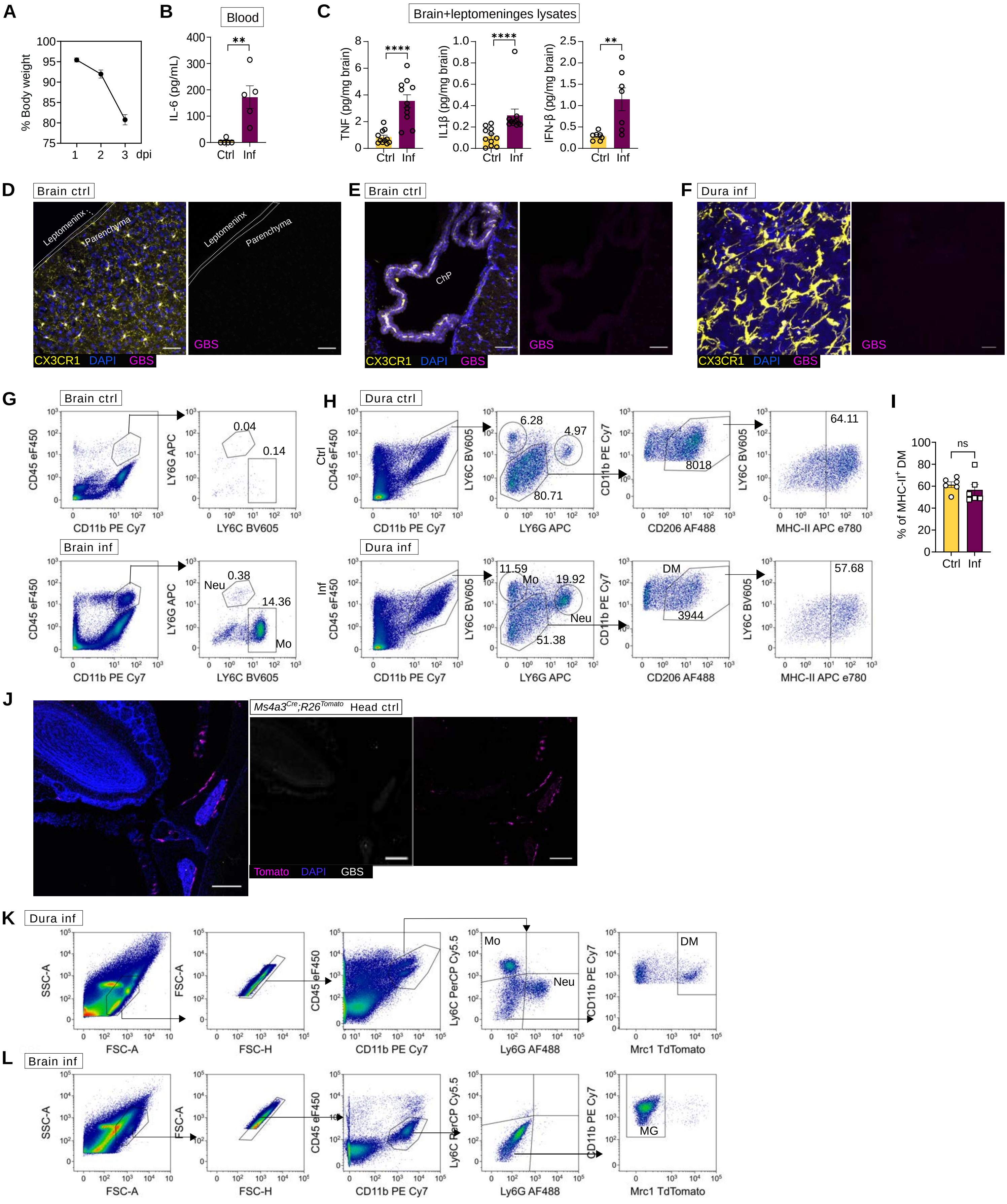
A
% Body weight
dpi
B
Blood
IL-6 (pg/mL)
Ctrl Inf
C
Brain+leptomeninges lysates
TNF (pg/mg brain)
IL1β (pg/mg brain)
IFN-β (pg/mg brain)
Ctrl Inf
D
Brain ctrl
Leptomeninx
Parenchyma
CX3CR1 DAPI GBS
E
Brain ctrl
ChP
F
Dura inf
G
Brain ctrl
Brain inf
CD45 eF450
CD11b PE Cy7
LY6G APC
LY6C BV605
Neu
Mo
H
Dura ctrl
Dura inf
Ctrl
Inf
CD206 AF488
MHC-II APC e780
DM
I
% of MHC-II+ DM
ns
J
Ms4a3Cre;R26Tomato Head ctrl
Tomato DAPI GBS
K
Dura inf
L
Brain inf
SSC-A
FSC-A
FSC-H
Ly6C PerCP Cy5.5
Ly6G AF488
Mrc1 TdTomato
MG

**Fig. S1: Hematogenous GBS infection highlights the meninges as a reactive CNS barrier**

**Fig. S1: A**: Weight loss curve in the first three days of the infection, for 1 dpi n=20, for 2 dpi n=23 and for 3 dpi n=8.

**B**: IL-6 in plasma 2 dpi, n=5. Statistics by Mann-Whitney U-test. Data represents 2 independent experiments**.** $^{**}p < 0.01$

**C:** TNF, IL-1β and IFN-β ELISA from homogenized brains. Measured values are normalized with the brain mass in mg. For IFN-β n=7, TNF and IL-1β n=11. Statistics by Mann-Whitney U-test. Data represent at least three independent experiments. $^{**}p < 0.01$, $p^{****} < 0.0001$

**D-E:** Representative maximal intensity projection of sagittal brain slices of control $Cx3cr1^{gfp/+}$ mice including leptomeningeal area and parenchyma (**D**) or choroid plexus (**E**) stained with GBS-AF633 (magenta) and DAPI (blue). Scale bar = 50 µm.

**F:** Representative maximal intensity projection of extra-sinusoidal region of whole mount control dura from $Cx3cr1^{gfp/+}$ mouse, stained with GBS-AF633 (magenta) and DAPI (blue). Scale bar = 20 µm.

**G-H**: Representative flow cytometry plots of percentage of monocytes and neutrophils in control and infected (**G**) brain or (**H**) dura. In **G**, numbers indicate percentages of the population, out of all myeloid cells. In **H**, numbers represent percentages of monocytes (Mo) and neutrophils (Neu), numbers of DM, and percentages of MHC-II positive DM.

**I:** Comparison of % of MHC-II positive DM in control and infected mice, n= 6, from at least two independent experiments. Statistics by unpaired t-test. Ns = non significant.

**J:** Representative image of a coronal slice of control $Ms4a3^{Cre};R26^{Tomato}$ mouse, stained with DAPI (blue), and GBS-AF633 (white). Scale bar = 200 µm.

**K-L:** Representative flow cytometry gating used for sorting DM (**L**) and MG (**M**) from $Mrc1^{CreERT2};R26^{Tom}$ mice for bulk RNA sequencing.

### Fig. S2

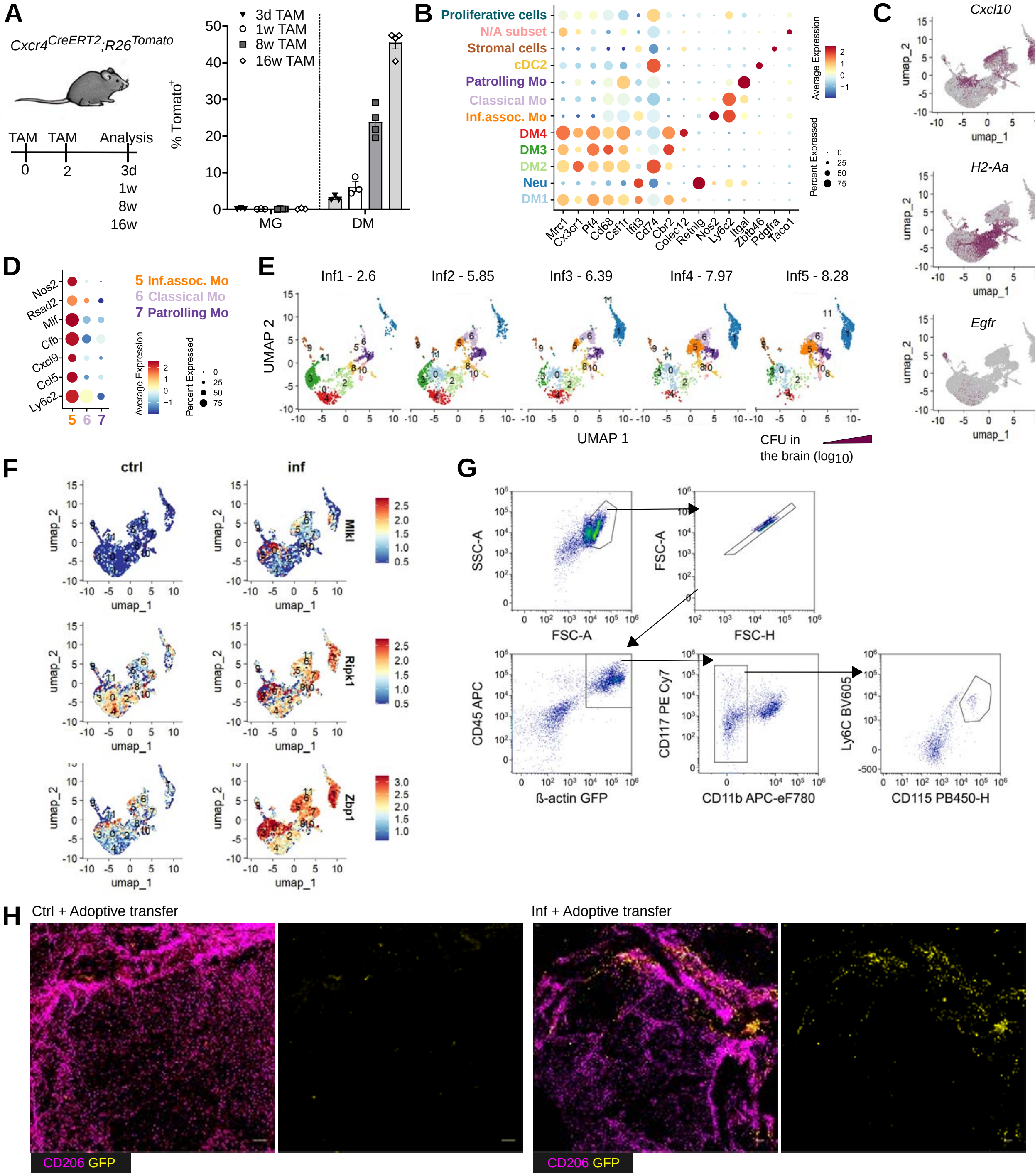

A
$Cxcr4^{CreERT2}$;$R26^{Tomato}$
TAM TAM Analysis
0 2 3d 1w 8w 16w
3d TAM
1w TAM
8w TAM
16w TAM
% Tomato+
MG
DM
B
Proliferative cells
N/A subset
Stromal cells
cDC2
Patrolling Mo
Classical Mo
Inf.assoc. Mo
DM4
DM3
DM2
Neu
DM1
Mrc1 Cx3cr1 Pf4 Cd68 Csf1r Ifit3 Cd74 Cbr2 Colec12 Retnlg Nos2 Ly6c2 Itgal Zbtb46 Pdgfra Taco1
Average Expression
Percent Expressed
C
Cxcl10
Oasl1
H2-Aa
Lyve1
Egfr
Mrc1
umap_1
umap_2
D
Nos2 Rsad2 Mif Cfb Cxcl9 Ccl5 Ly6c2
5 Inf.assoc. Mo
6 Classical Mo
7 Patrolling Mo
E
Inf1 - 2.6
Inf2 - 5.85
Inf3 - 6.39
Inf4 - 7.97
Inf5 - 8.28
UMAP 1
UMAP 2
CFU in the brain (log10)
F
ctrl
inf
Mlkl
Ripk1
Zbp1
G
SSC-A
FSC-A
FSC-H
CD45 APC
ß-actin GFP
CD117 PE Cy7
CD11b APC-eF780
Ly6C BV605
CD115 PB450-H
H
Ctrl + Adoptive transfer
Inf + Adoptive transfer
CD206 GFP
CD206 GFP

**Fig. S2: Loss of dural macrophages triggers compensatory monocyte differentiation**

**A:** *Cxcr4$^{CreERT2}$;R26$^{Tomato}$* mice were induced for 3 days, 1,8 or 16 weeks, recombination in DM and MG was measured as a percentage of Tomato positive $CD45^{hi}CD11b^{+}Ly6G^{neg}CD206^{+}$ DM in dura or $CD45^{lo}CD11b^{+}Ly6C^{neg}Ly6G^{neg}CD206^{neg}$ MG in parenchyma. n=3-4.

**B:** Dotplot representing selected discriminatory genes between different clusters. Color represents the mean expression of the gene and the dot size represents the percentage of the cells expressing the gene in each cluster.

**C:** Featureplot representing selected genes upregulated in each of the DM clusters.

**D:** Dotplot representing some inflammation-associated genes expressed in infection-associated monocytes, in comparison to classical and patrolling monocytes. Color represents the mean expression of the gene and the dot size represents the percentage of the cells expressing the gene in each cluster.

**E:** UMAP clustering of individual infected mice from least bacterial burden in the brain (inf1) to most (inf5) show burden-related inf-assoc. monocyte (cluster 5) and neutrophil (cluster 1) influx, in parallel with DM disappearance (clusters 0, 2, 3, 4).

**F:** Feature plots of canonical necroptosis genes expressed in control and infected condition.

**G:** Gating strategy for sorting MP from bone marrow of *β-actin-GFP* mice for adoptive transfer.

**H:** Representative image of whole mount dura, either control and infected, with donor-derived cells marked in yellow. Scale bar = 200 µm.

### Fig. S3

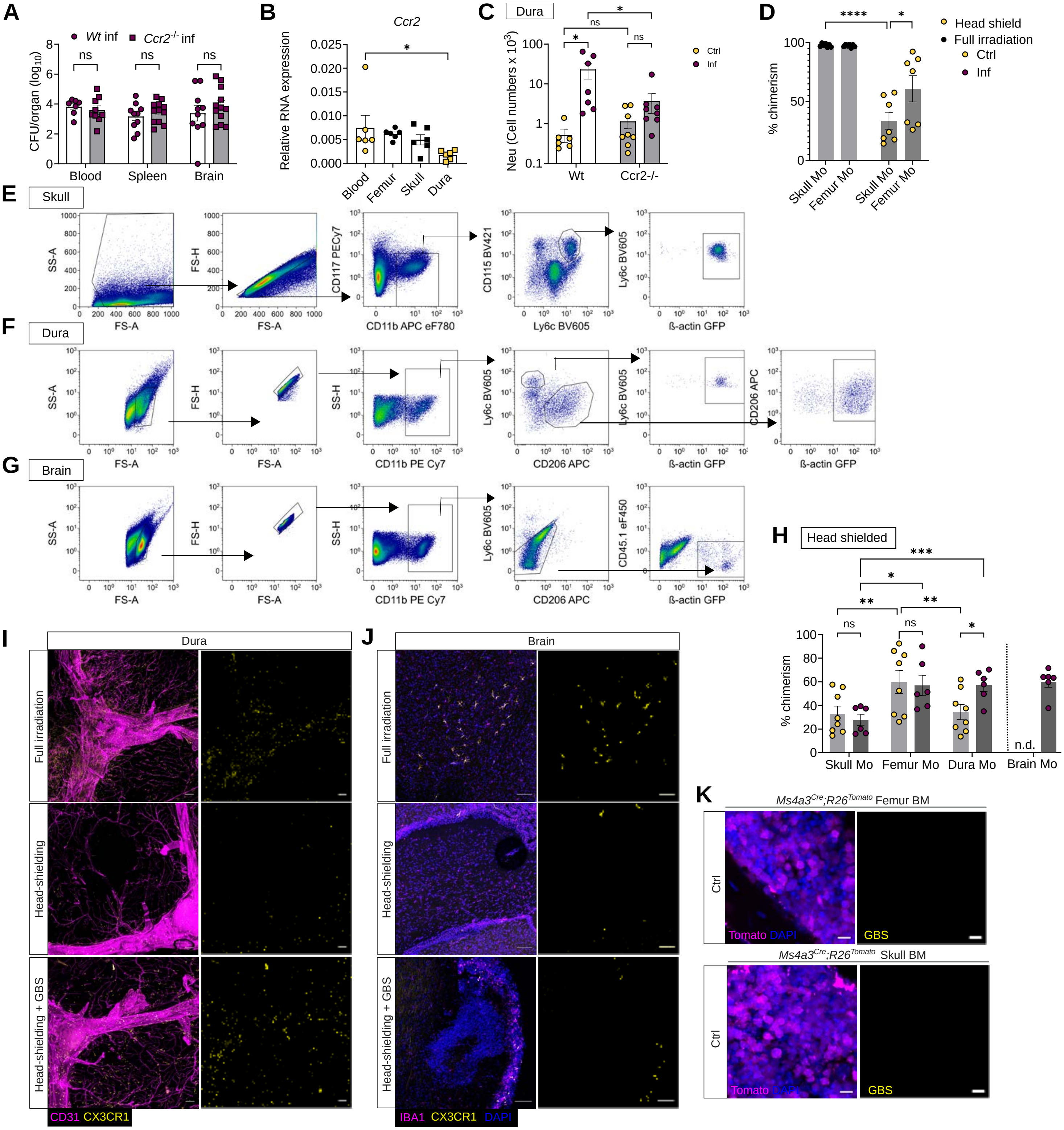

A
Wt inf
Ccr2-/- inf
CFU/organ (log10)
ns
Blood
Spleen
Brain
B
Ccr2
Relative RNA expression
Blood
Femur
Skull
Dura
C
Dura
Neu (Cell numbers x 10³)
Ctrl
Inf
Wt
Ccr2-/-
D
Head shield
Full irradiation
Ctrl
Inf
% chimerism
Skull Mo
Femur Mo
E
Skull
SS-A
FS-A
FS-H
CD117 PECy7
CD11b APC eF780
CD115 BV421
Ly6c BV605
ß-actin GFP
F
Dura
SS-H
CD11b PE Cy7
CD206 APC
G
Brain
CD45.1 eF450
H
Head shielded
Dura Mo
Brain Mo
n.d.
I
Dura
Full irradiation
Head-shielding
Head-shielding + GBS
CD31
CX3CR1
J
Brain
IBA1
DAPI
K
Ms4a3Cre;R26Tomato Femur BM
Ms4a3Cre;R26Tomato Skull BM
Ctrl
Tomato
GBS

**Fig. S3: Distinct monocyte sources support dural immunity in steady state and infection**

**A:** CFU counts from blood, spleen and brain of infected *Wt* or *Ccr2*$^{-/-}$ *mice*, n=7-12, statistics by multiple U-tests followed by Holm-Šídák, data represent at least three independent experiments. ns= not significant.

**B:** *Ccr2* expression measured by qPCR from monocytes sorted from blood, femur, skull and dura (n=6) of *Wt* mice. Statistics by one-way ANOVA and Bonferroni's multiple comparisons test. *p<0.05

**C:** Flow cytometry analysis of neutrophils in dura of control and infected *Wt* or *Ccr2*$^{-/-}$ mice. Statistics by two-way ANOVA and Tukey's multiple comparisons test (n=6-8). Data represent at least three independent experiments. *p<0.05

**D:** Percentage chimerism in head shielded or fully irradiated monocytes from skull or femur, measured by flow cytometry. Statistics by two-way ANOVA and Tukey's multiple comparisons test. N=7 from at least three independent experiments. *p<0.05, ****p<0.0001

**E-G**: Representative flow cytometry gating of donor-derived monocytes in skull (**E**), donor-derived monocytes and DM in dura (**F**) and donor-derived MG in brain (**G**) of fully irradiated mice.

**H:** Chimerism of monocytes in skull, femur, dura and brain of control and infected head-shielded chimeras. Two-way ANOVA followed by Tukey's multiple comparisons test, n=6-8, from at least three independent experiments. Brain monocytes were excluded from statistical analysis, due to low numbers of monocytes in steady state brains. *p< 0.05, **p<0.01,***p<0.001, ns = not significant, n.d. = not determined.

**I:** Representative images of whole-mounted dura from either fully irradiated and head shielded controls, or head shielded infected mice. Donors were *Cx3cr1*$^{+/gfp}$ mice. Donor derived cells are in yellow. Scale bar = 200 µm.

**J:** Representative images of brain sagittal sections from either fully irradiated and head shielded controls, or head shielded infected mice. Donors were *Cx3cr1*$^{+/gfp}$ mice. Donor derived cells are in yellow. Scale bar = 100 µm.

**K:** Femur and skull bone marrow from control *Ms4a3*$^{Cre}$*;R26*$^{Tomato}$ mice stained with GBS-AF633. Scale bar = 10 µm.

**Fig. S4**

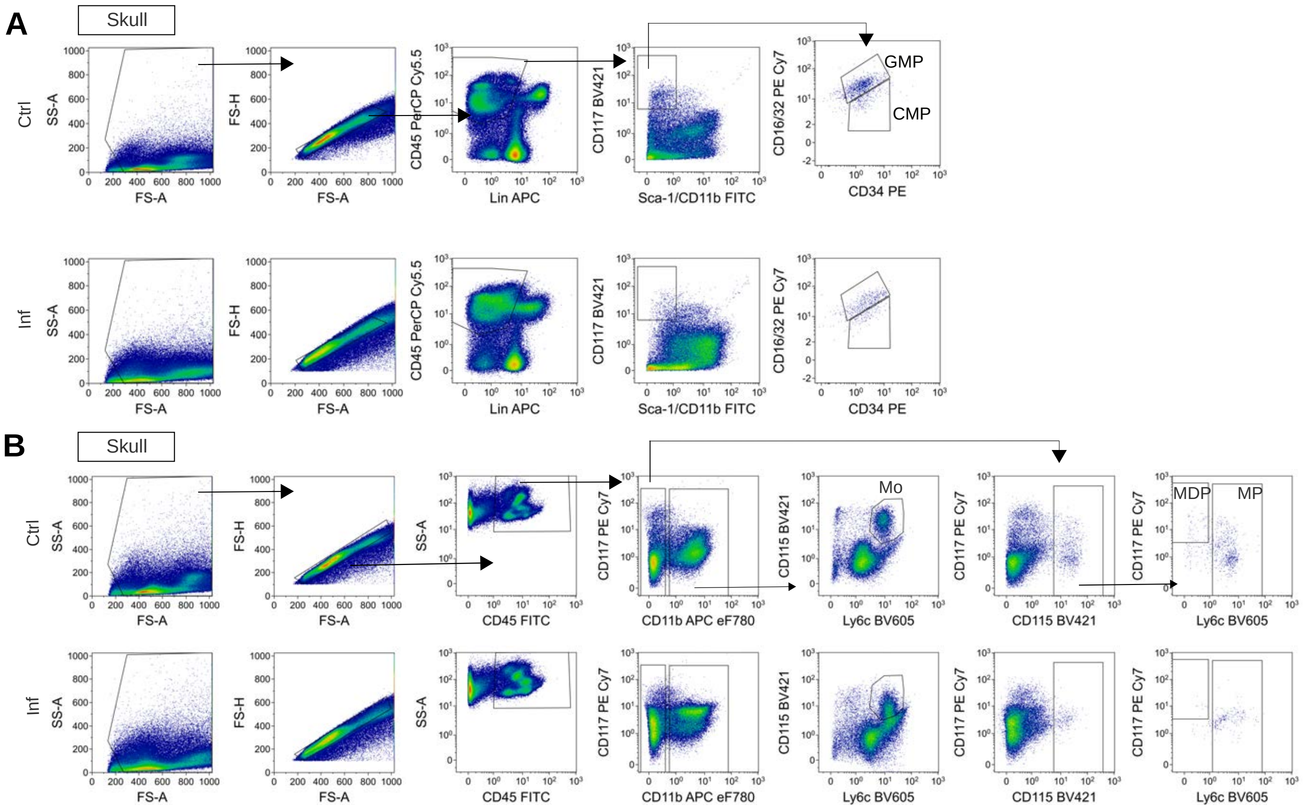
A
Skull
Ctrl
Inf
SS-A
FS-A
FS-H
CD45 PerCP Cy5.5
Lin APC
CD117 BV421
Sca-1/CD11b FITC
CD16/32 PE Cy7
CD34 PE
GMP
CMP
B
Skull
CD45 FITC
CD117 PE Cy7
CD11b APC eF780
CD115 BV421
Ly6c BV605
Mo
MDP
MP

**Fig. S4: Gating strategy**

**A:** Representative flow cytometry plots showing the gating strategy for CMP and GMP in control and infected skull bone marrow.

**B:** Representative flow cytometry plots showing the gating strategy for monocytes (Mo), MDP and monocyte progenitors (MP) in control and infected skull bone marrow.

#### Fig. S5

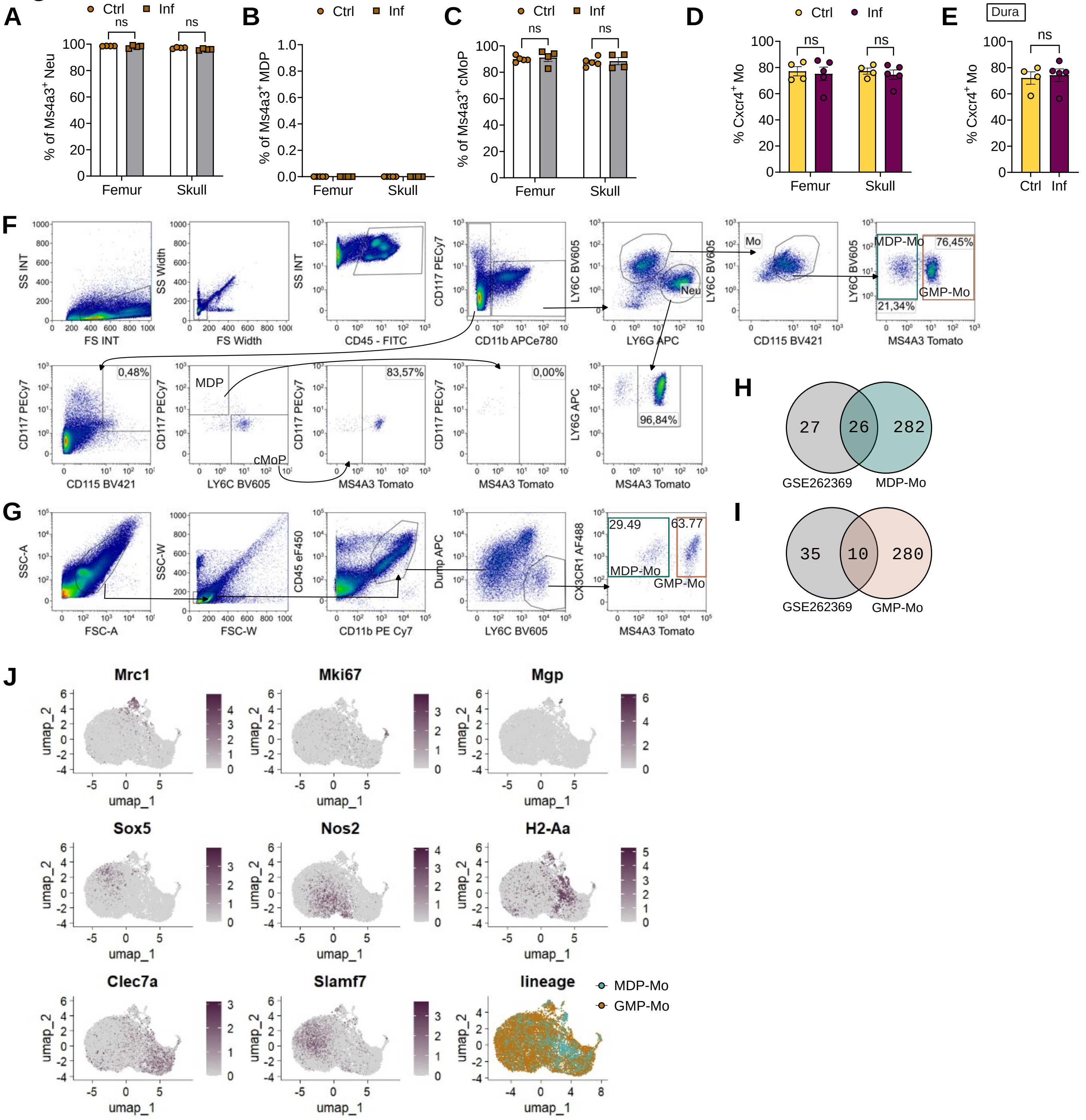
A
Ctrl
Inf
ns
% of Ms4a3+ Neu
Femur
Skull
B
% of Ms4a3+ MDP
C
% of Ms4a3+ cMoP
D
% Cxcr4+ Mo
E
Dura
F
SS INT
FS INT
SS Width
FS Width
CD45 - FITC
CD117 PECy7
CD11b APCe780
LY6C BV605
LY6G APC
Neu
Mo
CD115 BV421
MDP-Mo
GMP-Mo
76,45%
21,34%
MS4A3 Tomato
0,48%
MDP
cMoP
83,57%
0,00%
96,84%
G
SSC-A
FSC-A
SSC-W
FSC-W
CD45 eF450
CD11b PE Cy7
Dump APC
CX3CR1 AF488
29.49
63.77
H
27
26
282
GSE262369
I
35
10
280
J
Mrc1
Mki67
Mgp
Sox5
Nos2
H2-Aa
Clec7a
Slamf7
lineage
umap_1
umap_2

**Fig. S5: GBS infection reprograms monocyte ontogeny toward MDP-derived lineages**

**A-C:** Percentages of Tomato-positive neutrophils (Neu) (**A**), MDP (**B**) and cMoP (**C**) in *Ms4a3$^{Cre}$;R26$^{Tomato}$* mice in skull and femur, in steady state or 2 dpi. n=4-5, from two independent experiments. Statistics by two-way ANOVA and Šídák's multiple comparisons test. ns=not significant

**D:** Percentages of Tomato-positive monocytes (Mo) from skull or femur of *Cxcr4$^{CreERT2}$;R26$^{Tomato}$* mice in steady state or 2 dpi. n=4-5, from two independent experiments. Statistics by two-way ANOVA and Šídák's multiple comparisons test. ns=not significant

**E:** Percentages of Tomato-positive monocytes (Mo) from dura of *Cxcr4$^{CreERT2}$;R26$^{Tomato}$* mice in steady state or 2 dpi. n=4-5, from two independent experiments. Statistics by unpaired t-test. ns=not significant

**F-G:** Gating strategy for analysis in skull and femur (**F**) or dura (**G**) from *Ms4a3$^{Cre}$;R26$^{Tomato}$* mice. Plots represent infected skull (**F**) and infected dura (**G**).

**H:** Venn diagram comparing DEG from GSE262369 (MDP-Mo isolated from steady state bone marrow) and MDP-Mo DEG from pseudobulk anaylsis from infected dura. DEG are defined as padj<0.05 and $log_2$(fold change)>0.5.

**I:** Venn diagram comparing DEG from GSE262369 (GMP-Mo isolated from steady state bone marrow) and GMP-Mo DEG from pseudobulk anaylsis from infected dura. DEG are defined as padj<0.05 and $log_2$(fold change)>0.5.

**J:** Feature plots showing expression of genes specifically upregulated in certain cluster areas.

### Fig. S6

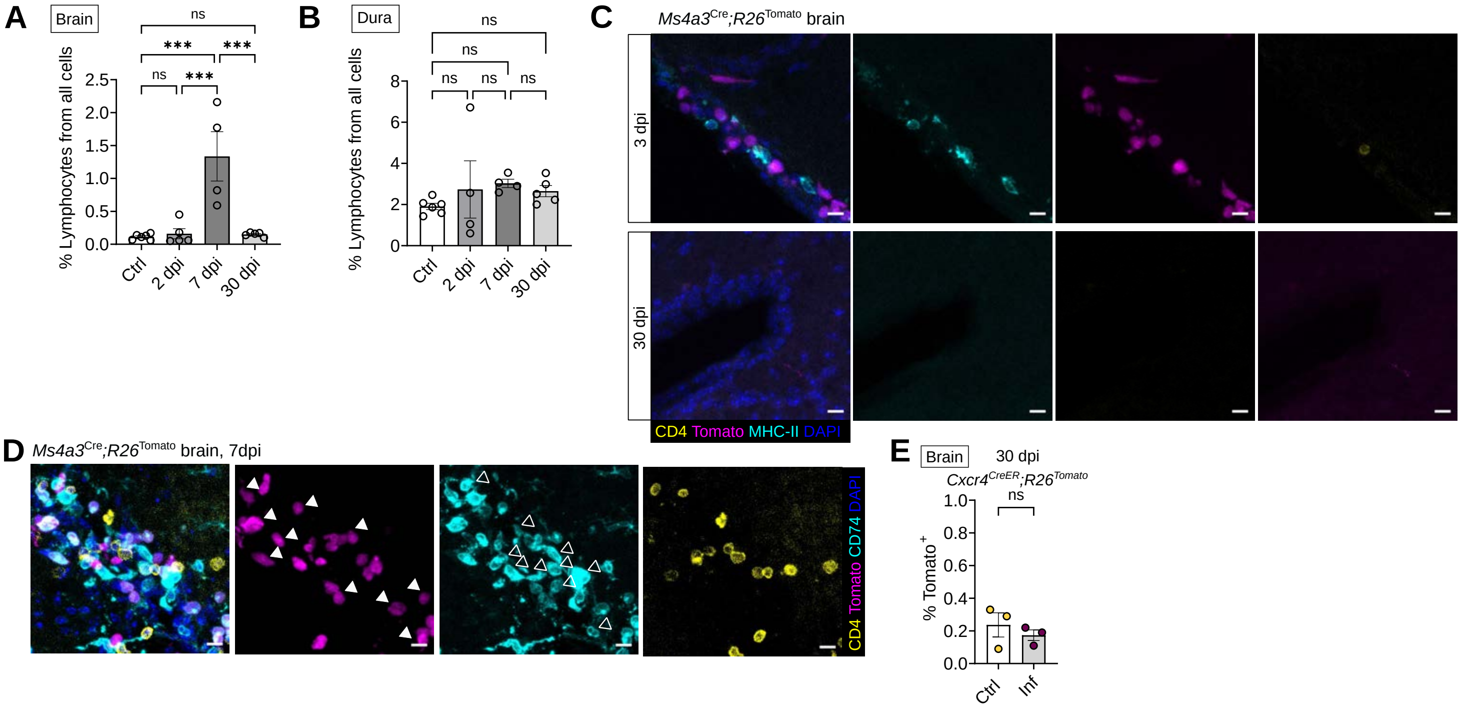

A
Brain
% Lymphocytes from all cells
0.0
0.5
1.0
1.5
2.0
2.5
ns
***
***
ns
***
Ctrl
2 dpi
7 dpi
30 dpi
B
Dura
% Lymphocytes from all cells
0
2
4
6
8
ns
ns
ns
ns
ns
Ctrl
2 dpi
7 dpi
30 dpi
C
Ms4a3Cre;R26Tomato brain
3 dpi
30 dpi
CD4 Tomato MHC-II DAPI
D
Ms4a3Cre;R26Tomato brain, 7dpi
CD4 Tomato CD74 DAPI
E
Brain
30 dpi
Cxcr4CreER;R26Tomato
ns
% Tomato+
0.0
0.2
0.4
0.6
0.8
1.0
Ctrl
Inf

**Fig. S6: Post-infection monocyte ontogeny diverges between dura and brain**

**A-B**: Flow cytometry analysis of the percentage of $CD11b^{neg}$, $CD45^{+}$ lymphocytes in brain (**A**) and dura (**B**). Statistics by one-way ANOVA, followed by Tukey's multiple comparisons test. N=4-5, data from two independent experiments. Ns= non significant, ***p < 0.001.

**C:** Representative microscopy images showing brain at 3 dpi and 30 dpi, from $Ms4a3^{Cre};R26^{Tomato}$ mice. Positions represented are those where many CD4 T-cells are found at 7 dpi. Scale bar = 10 µm.

**D:** Representative microscopy images from brain of $Ms4a3^{Cre};R26^{Tomato}$ mouse showing brain at 7 dpi. CD74 was used to discriminate between MDP-Mo and GMP-Mo, however, it is not a completely discriminatory marker. The full arrows point to $Ms4a3^{+}CD74^{+}$ cells, while the empty arrows point to cells that are $Ms4a3^{neg}CD74^{+}$, presumably MDP-Mo. Scale bar = 10 µm.

**E:** Percentages of Tomato-positive MG in brain at 30 dpi, from $Cxcr4^{CreERT2};R26^{Tomato}$ mice. Statistics by Student t-test, n=3.
